## Supplementary Figures for "A genome-wide CRISPR/Cas9 screen identifies genes that regulate the cellular uptake of α-synuclein fibrils by modulating heparan sulfate proteoglycans"

**Supplementary Figure 1 – Recombinant α-syn-PFFs characterization by electron microscopy**

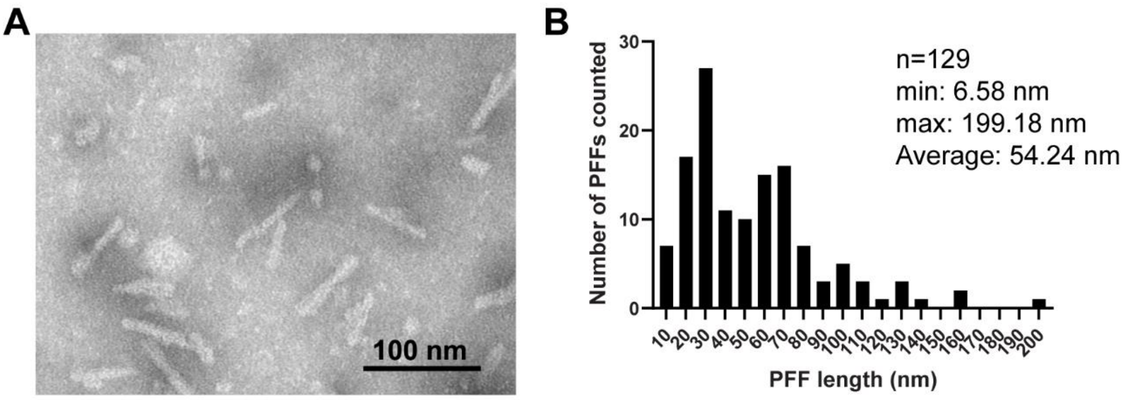


**(A)** Recombinant α-syn-PFFs purified from E. coli were stained with 2 % Uranyl Acetate and analyzed in a Tecnai 12 120 kV TEM microscope. **(B)** The length of PFFs was measured using ImageJ and the length distribution was obtained using MatLab.

**Supplementary Figure 2 – Cell size is a confounding factor in FACS-based genome-wide CRISPR screening**

The fact that most “inhibitors” of α-syn PFF accumulation were related to cell growth/cell cycle processes **(Figure 1C)** pointed us towards a possible cell size-related bias in our screen. This was confirmed by several lines of evidence. FACS analysis of the cellular pool from the screen treated with fluorescent α-syn PFF showed a significant correlation (R=0.764) between PFF content and the FSC value, that measures cell size **(Supplementary Figure 2B)**. This indicates that any treatment that will modify cell size is likely to affect total PFF content. In actively proliferating RPE-1 cells, this is likely to occur whenever cell cycle progression is affected. In our pooled genome-wide screen, this can occur upon disruption of cell cycle or cell growth related genes. Accordingly, disruption of *TP53* and *CDKN1A* tumor suppressor genes is known to lead to an increased cell proliferation due to defective G1/S checkpoint. This would result in smaller cells, which would in turn have a lower PFF content **(Figure 1B)**. Inversely, inactivation of oncogenes (*KIF11*, *MTBP*, *POLE2*, *TOP2A*,…) would result in slower cell-cycle progression (due to cell cycle blockade at the G2/M checkpoint for instance (*TOP2A*, *POLE2*)), and an overall increased cell size that would correlate with increased PFF content, and thus enrichment of target sgRNAs in the “High PFF” population. Beyond the inactivation of specific cell cycle-related genes, it was previously shown that double-strand breaks (DSBs) induced by the CRISPR/Cas9 system itself leads to cell cycle arrest in a p53-dependent manner, independent of the target locus. This likely explains the enrichment for non-targeting sgRNAs from the library (*LacZ*, *EGFP*, *Luciferase*, and Y chromosome-targeting in the XX chromosome-bearing RPE-1 cell line) in the “Low PFF” population, since such Cas9/sgRNA complexes are incompetent for DSB formation in the absence of the target DNA sequence **(Supplementary Figure 2C)**. The absence of the DSB-associated DNA damage response in these cells compared to all other cells from the pooled genome-edited population likely allows them to cycle faster relative to other cells, correlating with a reduced size and lower PFF content. This was experimentally confirmed by comparing a cell line expressing *LacZ* sgRNA/Cas9 complexes to the pooled genome-edited population **(Supplementary Figure 2A)**. *LacZ* sgRNA-bearing cells had lower FSC value, and upon PFF treatment, their median PFF fluorescence value was well below the lower quartile of the pooled genome-edited population, in line with their enrichment in the “Low PFF” population upon FACS sorting based solely on PFF fluorescence. The PFF fluorescence/FSC ratio however shows that *LacZ* sgRNA-bearing cells have PFF amount relative to cell size that is comparable with the pooled population. Together, this data demonstrates the importance of normalizing fluorescent signals to cell size in FACS-based genome-wide screens.

**Supplementary Figure 2 (continued)**

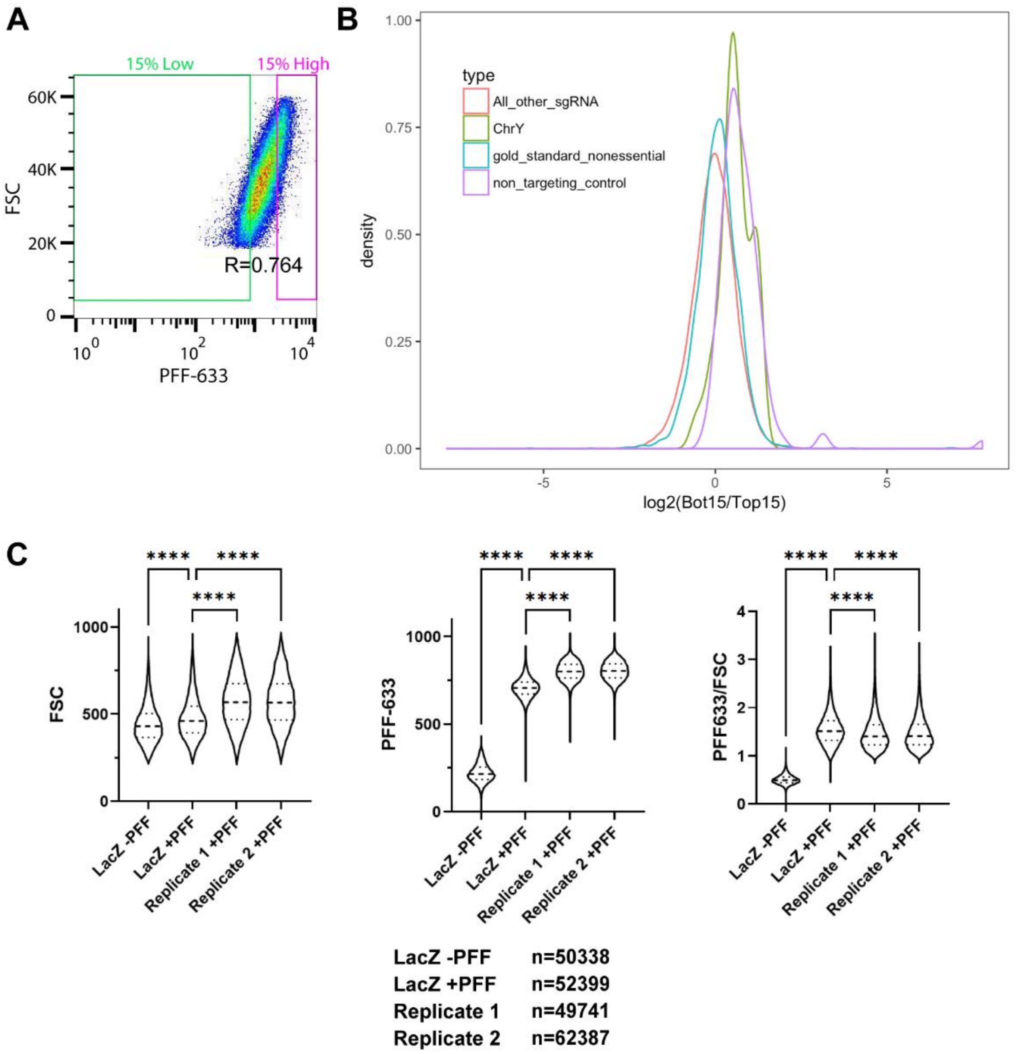


**(A)** Dot plot of FSC values as a function of PFF-A633 fluorescence for cells transduced with the genome-wide CRISPR library. Linear regression was performed and the correlation coefficient (R) was calculated, showing significant correlation between cell size and PFF-A633 fluorescence. **(B)** Densities of indicated sgRNA groups enrichment in the Low PFF population versus the High PFF population, showing that sgRNAs that don't result in double strand break (non-targeting control, and ChrY targeting sgRNAs in RPE-1 cells from female origin) are enriched in the Low PFF population, most probably as a result of a lower cell size (see **(C)**). **(C)** Violin plots showing FSC (left), PFF-A633 (middle) and PFF-A633/FSC (right) values distribution for n>49500 cells per condition from samples analyzed by FACS at the end of the genome-wide CRISPR screen. A control population transduced with a non-targeting control sgRNA (*LacZ*) was untreated or treated with 15 nM PFF-A633 at the same time as the two replicate populations from the actual genome-wide CRISPR library-transduced cells. The dashed lines and dotted lines indicate the median values and quartiles respectively. Statistical test: one-way ANOVA ; **** p<0.0001.

**Supplementary Figure 3 – Hits selection, validation of screen validation methodology, and α-syn-PFFs uptake quantification in polyclonal KO cells for validated hits**

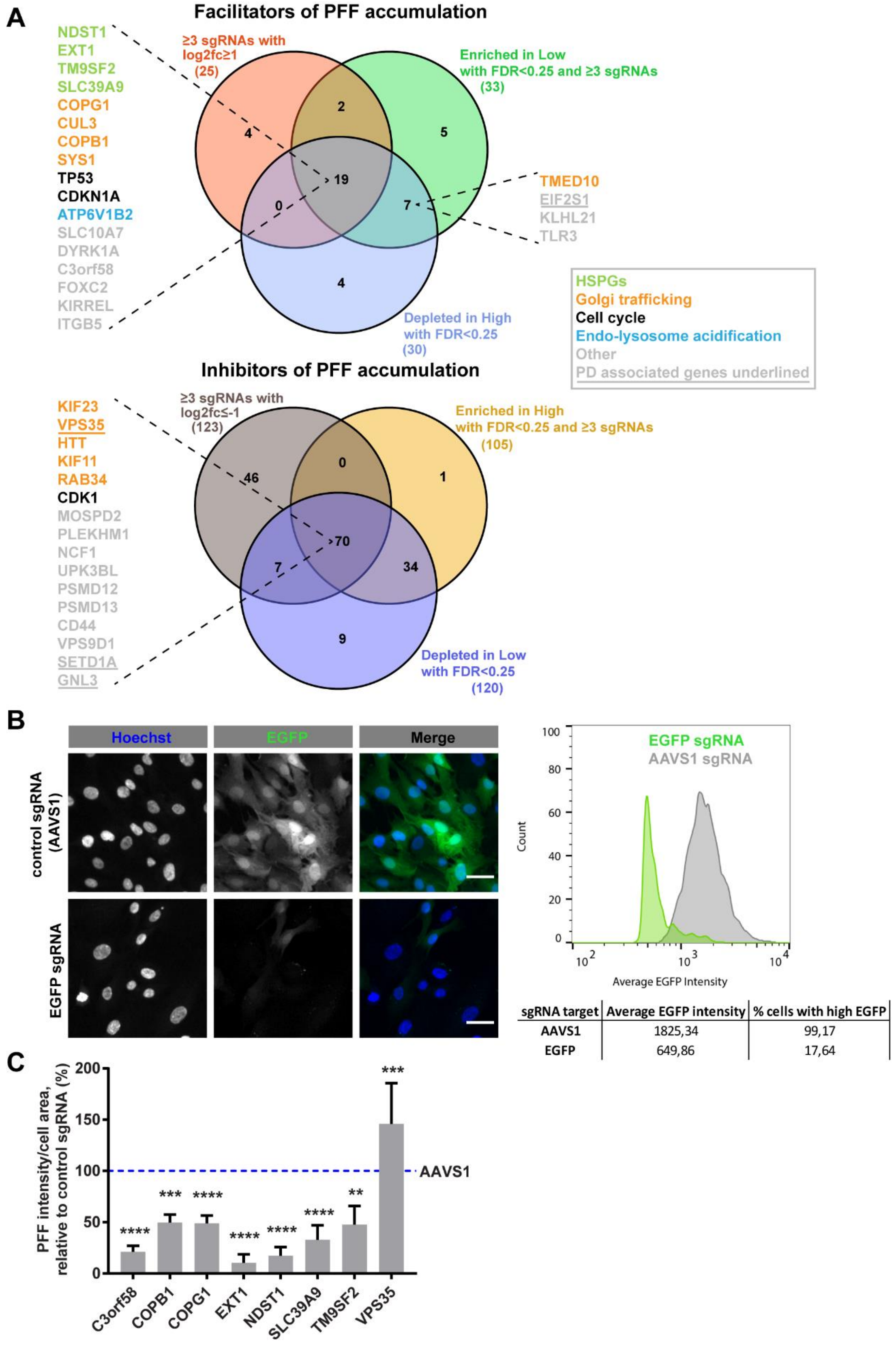


**Supplementary Figure 3 (continued)**

**(A)** Based on MAGeCK sgRNA enrichment analysis (see Supplementary Tables S1, S2 and S3) between the low PFF vs high PFF populations, and the high PFF vs low PFF populations counter analysis, we selected hits to further validate that met the following criteria: 1) 3 or 4 sgRNAs with log2 fold-change ≥1 (for facilitators of PFF uptake) or ≤-1 (for inhibitors of PFF uptake) ; 2) enrichment in the low or high PFF population with FDR <0.25 and 3 or 4 "good sgRNAs" ; 3) depletion in the high or low PFF population with FDR <0.25 and 3 or 4 "good sgRNAs". Further filtering was done to remove most genes with ontologies related to cell growth or cell cycle (based on results from GOrilla analysis). Some hits were also cherry-picked with relaxed criteria, because of their location in genetic loci associated with PD (*EIF2S1*), or their good ranking despite some sgRNAs having magnitude of log2 fold-change not reaching the ≥1 cut-off (*TMED10*, *KLHL21*, *TLR3*) **(B)** RPE-1 cells stably expressing Cas9 and EGFP were transfected with a control AAVS1-targeting or EGFP-targeting sgRNA for 3 days before fixation, nuclei staining with Hoechst 33342 and analysis using a CX7 high-content microscope. The average EGFP intensity per cell and % cells with high EGFP intensity were quantified (table on the right). The results of cellular analysis were exported as FCS files, and the FlowJo software was used for graphic representation of both cell populations (*AAVS1* sgRNA-treated in grey, *EGFP* sgRNA-treated in green). Scale bar: 30 µm. **(C)** The results of PFF intensity/cell area from Figure 2B (TKOv3 sgRNA) of the high-content microscopy-based validation are shown as a histogram for the indicated genes. Statistical test: one-way ANOVA; ** $p<0.01$, *** $p<0.001$, **** $p<0.0001$.

**Supplementary Figure 4 – Validation for cargo specificity**

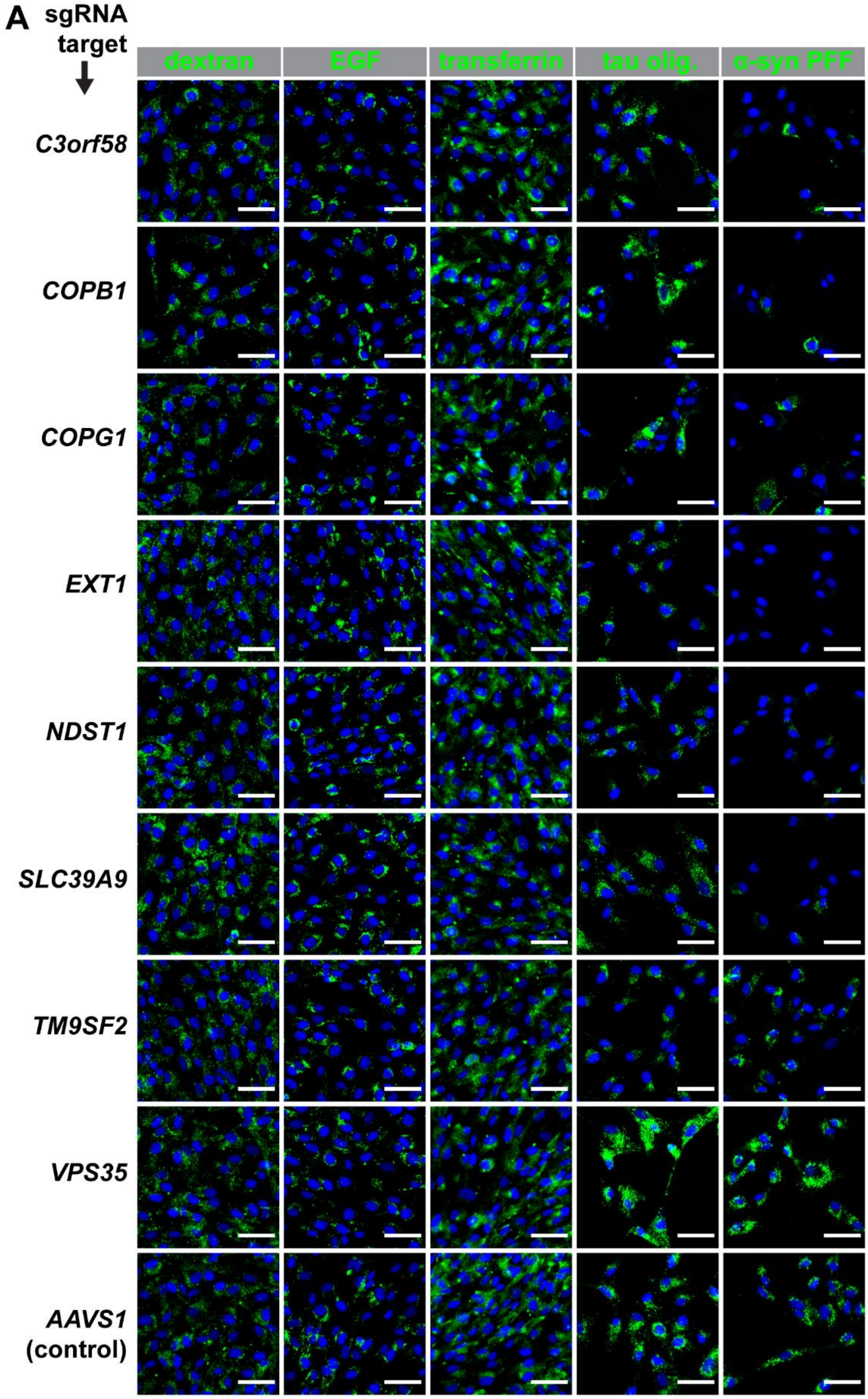

**Supplementary Figure 4 (continued)**

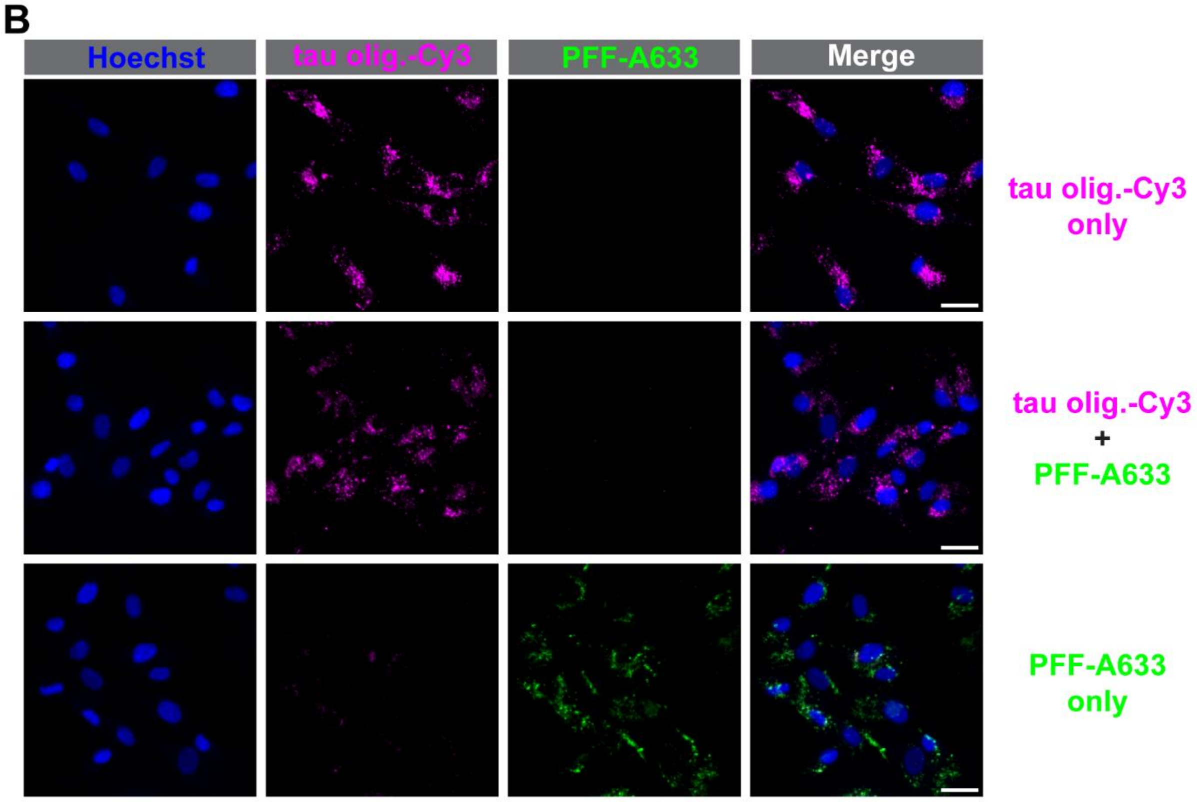


**(A)** RPE-1 cells stably expressing Cas9 nuclease were transfected in 96 wells plates with individual synthetic sgRNAs, and after 3 days, were subjected to 24 h uptake of the indicated cargoes (green): dextran-Oregon Green 488, EGF-A488, transferrin-A488, Tau oligomers-Cy3, and α-syn PFF-A633 as positive controls of sgRNAs' effects. Cells were then fixed, and nuclei stained with Hoechst 33342 (blue). Imaging was done using a CX7 high-content microscope. Quantifications are shown for each cargo in **Figure 3**. Scale bar: 30 µm. **(B)** WT RPE-1 cells were treated for 24 h with either 80 nM Tau oligomers-Cy3 (magenta), 60 nM α-syn-PFF-A633 (green) or both before fixation, nuclei staining with Hoechst (blue), and imaging using a CX7 high-content microscope. Scale bar: 30 µm.

Supplementary Figure 5 – The PAPS transporters SLC35B2 and SLC35B3 regulate α-syn PFF uptake

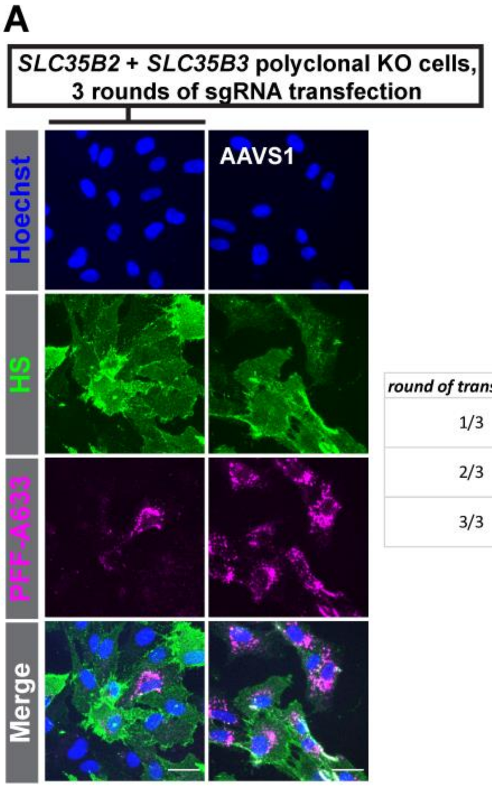


| round of transfection | sgRNAs used |
|---|---|
| 1/3 | SLC35B2_3 |
| | SLC35B3_1 |
| 2/3 | SLC35B2_3 |
| | SLC35B3_1 |
| 3/3 | SLC35B2_1 |
| | SLC35B3_2 |

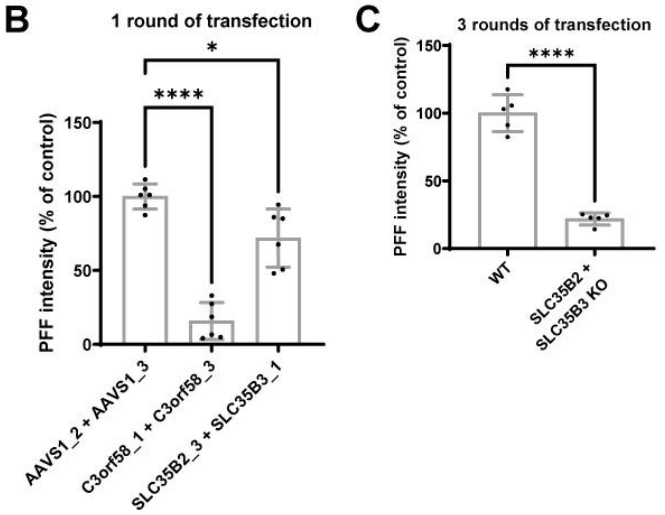

**(A)** RPE-1 cells expressing Cas9 were subjected to 3 rounds of transfection with the indicated sgRNAs combinations **(table)** against the redundant PAPS transporters genes *SLC35B2* and *SLC35B3.* Then, cells were treated for 24h with PFF-A633. After fixation, unpermeabilized cells were stained for HS (10e4 antibody, green) and nuclei were stained with Hoechst (blue). PFF are in magenta. Imaging and quantification were done using a CX7 high-content microscope. Quantified in **(C)**. Scale bar: 30 µm. **(B)** Cells were transfected for 3 days with the indicated sgRNAs. Then, 24h PFF-A633 uptake was done before staining as in **(A)** and quantification. Bar graph showing mean ±SD of PFF signal intensity relative to negative control (parental RPE-1 Cas9 cells) and positive control of decreased PFF uptake (*C3orf58*). A 25% decrease in PFF uptake was observed after CRISPR-mediated disruption of both PAPS transporter genes simultaneously, after one round of transfection. **(C)** shows the quantification of the experiment in **(A)**, done after three rounds of transfections. An 80% decrease in PFF uptake was observed, presumably due to a higher proportion of cells having both genes disrupted in the population compared to a single round of transfection.

### Supplementary Figure 6 – sequences of monoclonal CRISPR-invalidated lines used

**A**

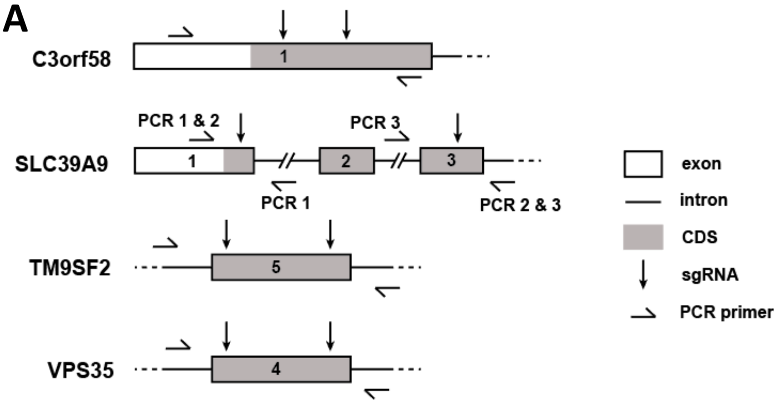


**B** *C3orf58*$^{-/-}$ clone 3

```
allele 1        MWRLVPPKLGRLSRSLKLAALGSLLVLMVLHSPSLLASWQRNELTPSGAQAPRL-AARAG    59
wt              MWRLVPPKLGRLSRSLKLAALGSLLVLMVLHSPSLLASWQRNELTDRRFLQLNKCPACFG    60
allele 2        MWRLVPPKLGRLSRSLKLAALGSLLVLMVLHSPSLLASWQRNELTRPALPAAQ*VPGVLR    59
                *********************************************       .   .

allele 1        AARPEHLQAGHRPA----PLRPAAGHAPDR--------VRAPQRRRASAHARGGGGLV--    105
wt              TSWCRRFLNGQVVFEAWGRLRLLDFLNVKNVYFAQYGEPREGGRRRVVLKRLGSQRELAQ    120
allele 2        ----------------------------------------------------------HE    61

allele 1        -----------GPGALPLAAPSRPPGAPLRGDQGLGQLPASQPQGLGAHAAAADPGLQPR    154
wt              LDQSICKRATGRPRCDLLQAMPRTEFARLNGDVRLL--TPEAVEGWSDLVHCPSQRLLD-    177
allele 2        LVPPLPQRAGGRPRCDLLQAMPRTEFARLNGDVRLL--TPEAVEGWSDLVHCPSQRLLD-    118
                    * .  * *  *   * *.**  *     .  :* .  . . .  *

allele 1        AAGATEFSV**RLAICKVSWSLWKNGGCKLCW-RR---TVE-LL*CAMGKTS*PRLAING    205
wt              -----------RL---VRRYAETKDSGSFLLRNLKDSERMQLLLTLAFNPEPLVLQSFPS    223
allele 2        -----------RL---VRRYAETKDSGSFLLRNLKDSERMQLLLTLAFNPEPLVLQSFPS    164
                   **      ::  *:.*. *    :    :: **  *:.       :: .

allele 1        NSRTAYKQ*L*ICTLPPGRQL*---QFCSWS*RWEGNHCGC*KCFGC*QKIN*TK*T*KL    253
wt              DEGWPFAKYLGACGRMVAVNYVGEELWSYFNAPWEKRVDLAWQLMEIAEQLTNNDFEFAL    283
allele 2        DEGWPFAKYLGACGRMVAVNYVGEELWSYFNAPWEKRVDLAWQLMEIAEQLTNNDFEFAL    224
                :.   : : *  *    . :      :. :.  ** .   . : :   :::. ..    *

allele 1        GC--MV*KQV**L**GGLLIIFKRNSLCSCHCGPQLLCCLSEPLIQTCHLAWHFWRTPS*    305
wt              YLLDVSFDNFAVGPRDGKVIIVDAEN---------VLVADKRLIRQNKPENWDVWYESKF    334
allele 2        YLLDVSFDNFAVGPRDGKVIIVDAEN---------VLVADKRLIRQNKPENWDVWYESKF    275
                    :  .:.     .* :**.. :.         :* . .. : *.    *..*   .

allele 1        ST--K*NCQRWPARGL---------AG*VCQP---KEALWQIPGCKRTA*IPSTIK*QRE    347
wt              DDCDKEACLSFSKEILCARATVDHNYYAVCQNLLSRHATWRGTSGGLLHDPPSEIAKDGR    394
allele 2        DDCDKEACLSFSKEILCARATVDHNYYAVCQNLLSRHATWRGTSGGLLHDPPSEIAKDGR    335
                .   *  *  :  . *            ***    :.* *:  .       ** *  : .

allele 1        V------------------------------------   348
wt              LEALLDECANPKKRYGRFQAAKELREYLAQLSNNVR*   430
allele 2        LEALLDECANPKKRYGRFQAAKELREYLAQLSNNVR*   371
                :
```

**Supplementary Figure 6 (continued)**

### C *SLC39A9*$^{-/-}$ clone 10

```
allele 1    MDDFISISLLSLAMLVGCYVAGII-LAVNFSEERLKLVTVLGAGLLCGTALAVIVPEGVH    59
wt          MDDFISISLLSLAMLVGCYVAGIIPLAVNFSEERLKLVTVLGAGLLCGTALAVIVPEGVH    60
allele 2    MDDFISISLLSLAMLVGCYVAGII------------------------------------    24
            ************************

allele 1    ALYEDILEGKHHQASETHNVIASDKAAEKSVVHEHEHSHDHTQLHAYIGVSCCWWTRLVT    119
wt          ALYEDILEGKHHQASETHNVIASDKAAEKSVVHEHEHSHDHTQLHAYIGVSLVLGFVFM-    119
allele 2    ---------------------------------------------------LLGFVFM-    31
                                                                     ::

allele 1    PMCILLTIQKQQGLA--IPKSPPRWVWL------SMLQLMVLLWEQQHLLHRPV-----S    166
wt          -----LLVD-QIGNSHVHSTDDPEAARSSNSKITTTLGLVVHAAADGVALGAAASTSQTS    173
allele 2    -----LLVD-QIGNSHVHSTDDPEAARSSNSKITTTLGLVVHAAADGVALGAAASTSQTS    85
                 * :: * * :    .. *. .        : * *:*    :   *   .     *

allele 1    S*LCLWQSCYIRHQLLLDWFPS*CMLA*SGIESESTCWSLHWQHQLCPW*HT---*D*VR    217
wt          VQLIVFVAIML-HKA-PAAFGLVSFLMHAGLERNRIRKHLLVFALAAPVMSMVTYLGLSK    231
allele 2    VQLIVFVAIML-HKA-PAAFGLVSFLMHAGLERNRIRKHLLVFALAAPVMSMVTYLGLSK    143
              * :: :  : *:     *   .:*  :*:* :     *      .*        .  :

allele 1    AVKKPFQR*TPREWPCFSLPGHFFMLPQYMSSLRWAE*GTATSPMPREGEASAAWKWQPW    275
wt          SSKEALSEVNATGVAMLFSAGTFLYVATVHVLPEVGGIGHSHKPDATGGRGLSRL-----    286
allele 2    SSKEALSEVNATGVAMLFSAGTFLYVATVHVLPEVGGIGHSHKPDATGGRGLSRL-----    198
            : *: :.. .      :   * *: :       . .  * : .*    *.. :

allele 1    FWVASSLSSCQ*DTSI-------  290
wt          -EVAALVLGCLIPLILSVGHQH*  307
allele 2    -EVAALVLGCLIPLILSVGHQH*  219
              **: : .*     :
```

### D *TM9SF2*$^{-/-}$ clone 11

```
WT          MSARLPVLSPPRWPRLLLLSLLLLGAVPGPRRSGAFYLPGLAPVNFCDEEKKSDECKAEI    60
alleles 1+2 MSARLPVLSPPRWPRLLLLSLLLLGAVPGPRRSGAFYLPGLAPVNFCDEEKKSDECKAEI    60
            ************************************************************

WT          ELFVNRLDSVESVLPYEYTAFDFCQASEGKRPSENLGQVLFGERIEPSPYKFTFNKKETC    120
alleles 1+2 ELFVNRLDSVESVLPYEYTAFDFCQASEGKRPSENLGQVLFGERIEPSPYKFTFNKKETC    120
            ************************************************************

WT          KLVCTKTYHTEKAEDKQKLEFLKKSMLLNYQHHWIVDNMPVTWCYDVEDGQRFCNPGFPI    180
alleles 1+2 KLVCTKTYHTEKAEDKQKLEFLKKSMLLNYQHHWIVDNMPVLLVQISMKEIHFTSSTMLT    180
            *****************************************       .  :* .  :

WT          GCYITDKGHAKDACVISSDFHERDTFYIFNHVDIKIYYHVVETGSMGARLVAAKLEPKSF    240
alleles 1+2 SKYTI--------MLLKLGPWEQD*WL-------------------------LNLNRKAS    206
            . *           ::. .  *:* :                           :*: *:
```

Only the first 240 amino acids are shown.

### E *VPS35*$^{+/-}$ clone 21

```
allele_1    MPTTQQSPQDEQEKLLDEAIQAVKVQSFQMKRCLDKNKLMDALKHASNMLGELRTSMLSP    60
WT          MPTTQQSPQDEQEKLLDEAIQAVKVQSFQMKRCLDKNKLMDALKHASNMLGELRTSMLSP    60
allele_2    MPTTQQSPQDEQEKLLDEAIQAVKVQSFQMKRCLDKNKLMDALKHASNMLGELRTSMLSP    60
            ************************************************************

allele_1    KSYYELYMAISDELHYLEVYLTDEFAKGRKVADLYELVQYAGNIIPRLYLLITVGVVYVK    120
WT          KSYYELYMAISDELHYLEVYLTDEFAKGRKVADLYELVQYAGNIIPRLYLLITVGVVYVK    120
allele_2    KSYYELYMAISDELHSVVCNIP*GVC-------FFEITFFS---VPEISYLM-------K    102
            *************** :   :   ..       ::*:. ::   :*.:  *:       *
```

Only the first 120 amino acids are shown. Allele 1 was identical to WT.

**(A)** Schematic representation of knock-out (KO) and PCR-screening strategies for the indicated genes in human cells. For all genes, allelic sequences from the indicated CRISPR-edited RPE-1 cell clones were PCR-amplified using Q5 polymerase and the indicated oligonucleotides. For *C3orf58* **(B)** and *SLC39A9* **(C)**, all PCR products were cloned using a PCR cloning kit, and 10 plasmids from individual bacterial clones were sequenced. For *TM9SF2* **(D)** and *VPS35* **(E)**, the PCR products were directly subjected to Sanger sequencing, and the resulting trace files were analyzed using ICE analysis (Synthego) against the WT sequences. The identified allelic sequences are shown and aligned to the reference WT sequence.

**Supplementary Figure 7 – Heparan sulfate staining in monoclonal CRISPR KO and/or rescue cells for *C3orf58*, *SLC39A9*, *TM9SF2* and *VPS35* & PFF uptake in monoclonal *TM9SF2* -/- and *VPS35* +/- cells**

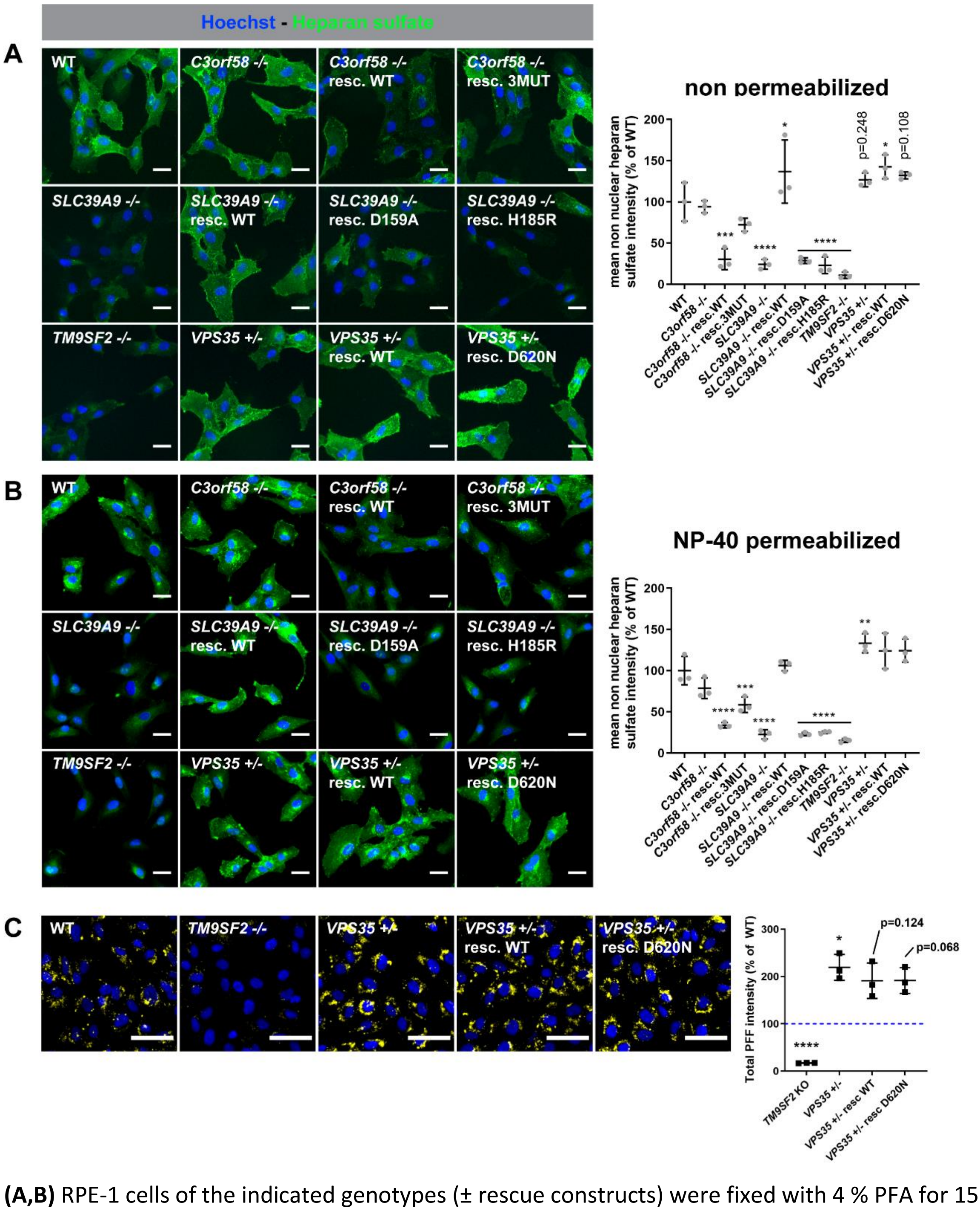


**(A,B)** RPE-1 cells of the indicated genotypes (± rescue constructs) were fixed with 4 % PFA for 15 min, then processed for immunostaining against Heparan sulfate (antibody clone 10e4, green) either in non-permeabilized conditions **(A)** or following a 10 min permeabilization with 0.05% NP-40 **(B)** to label surface or total heparan sulfate respectively. Nuclei were stained with Hoechst 33342 (blue). Scale bar: 30 µm. Cells were then imaged with a CX7 high-content microscope and analyzed with the HCS Studio Cell Analyzer software. Nuclei in the Hoechst channel used for object identification, and the mean of total non-nuclear heparan sulfate signal quantified for each condition. Data in graphs are shown as mean ±s.d., n=3. **(C)** 24h uptake of 60 nM PFF-A633 (yellow) in *TM9SF2* -/-or *VPS35* +/-

monoclonal KO RPE-1 cells, either in absence or presence of stable expression of the indicated C-terminally HA-tagged rescue contructs. Nuclei were stained with Hoechst (blue), and the mean PFF-A633 fluorescence intensity was quantified by high-content microscopy and normalized to WT RPE-1 cells (see graph, right panel). Statistical test: one-way ANOVA; * $p<0.05$, **** $p<0.0001$. Scale bar: 60 µm.

**Supplementary Figure 8 – Localization of SLC39A9 and C3orf58 in RPE-1 cells**

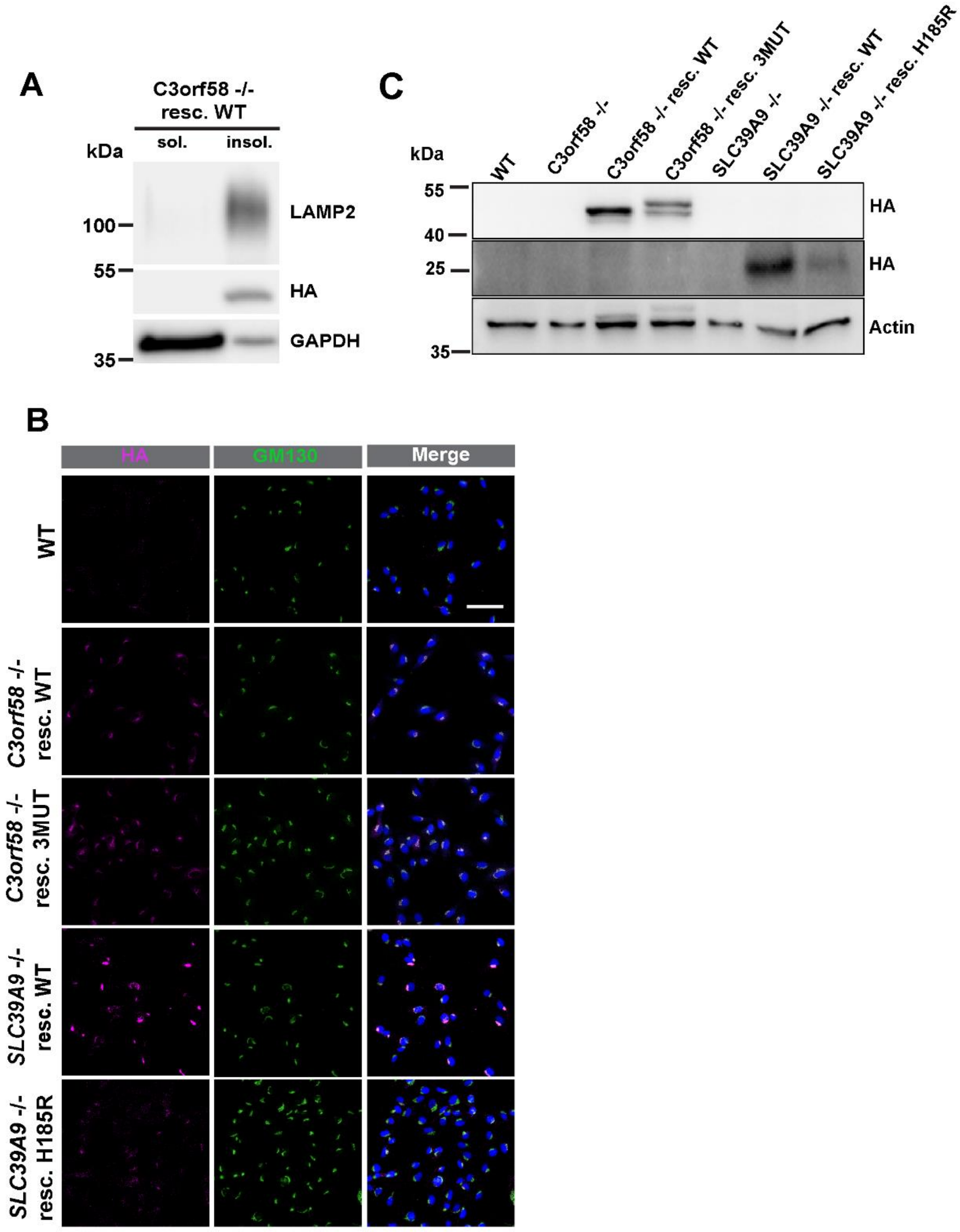


**(A)** Soluble and membrane associated proteins (soluble fraction = sol.) from *C3orf58* -/- RPE-1 cells rescued with WT C3orf58-HA were separated from integral membrane proteins (insoluble fraction = insol.) by 0.1 M sodium carbonate (pH 11.5) extraction followed by ultracentrifugation at 100,000 g. Both fractions were analyzed by Western blot against GAPDH (soluble protein), LAMP2 (transmembrane protein), and HA showing that C3orf58 is an integral membrane protein. **(B)** The monoclonal KO cells for *SLC39A9* or *C3orf58* were rescued using lentiviral transduction to drive puromycin-selectable constitutive expression of C-terminally HA-tagged constructs encoding WT SLC39A9, H185R (Zn transporter mutant) SLC39A9, WT C3orf58 or triple mutant (putative kinase-dead) C3orf58 as indicated. These cells as well as WT RPE-1 cells were then plated in imaging-compatible plates, fixed, permeabilized with 0.15% Triton X-100, and processed for immunofluorescence using anti-HA (magenta) and anti-GM130 (Golgi marker, green) antibodies. Nuclei were stained with Hoechst 33342 (blue), and images were acquired using a Nikon A1 laser scanning confocal microscope at 40x. Scale bar: 20 µm. **(C)** Western blot showing expression levels of HA-tagged C3orf58 WT and 3MUT, SLC39A9 and H185R Mut constructs in rescued monoclonal RPE-1 cells.

**Supplementary Figure 9 – SLC39A9 modulates Zinc homeostasis in the Golgi**

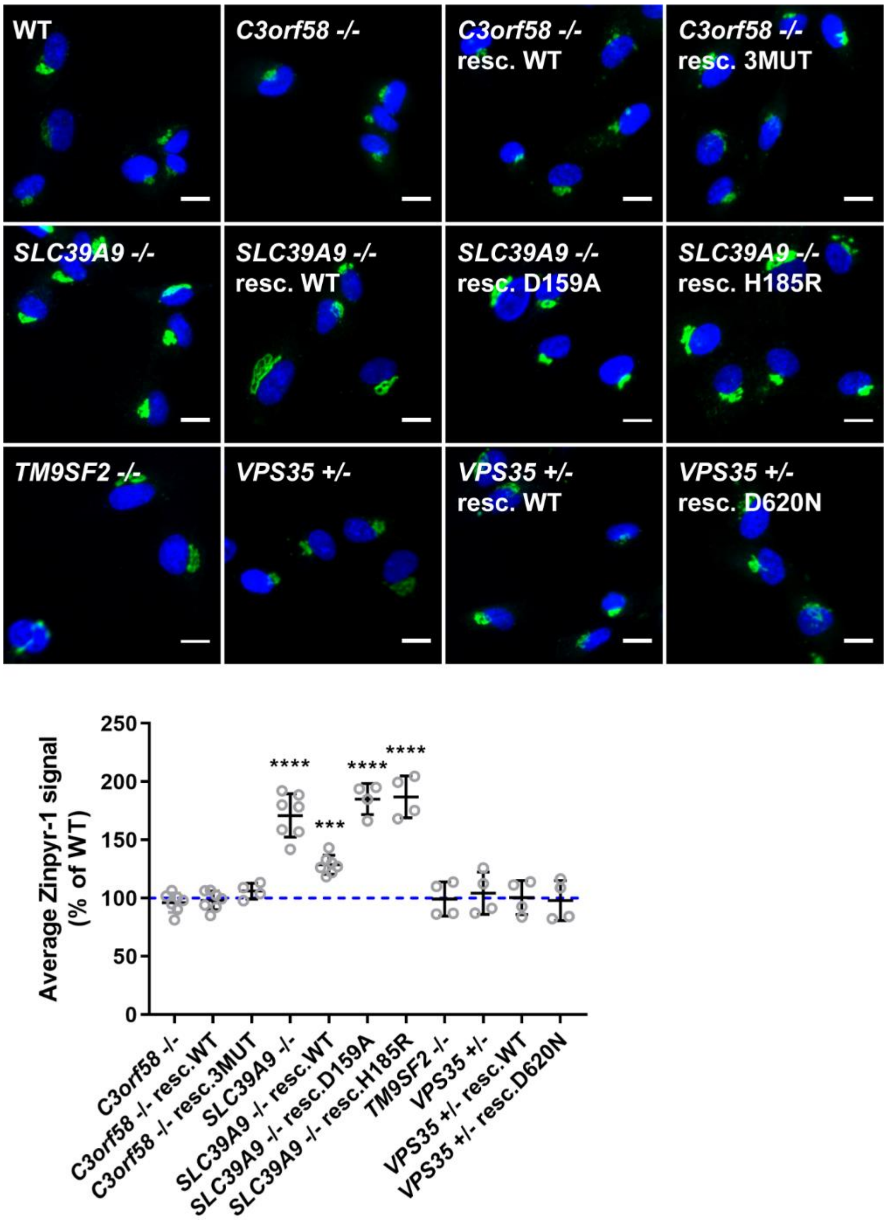


RPE-1 cells of the indicated genotypes were seeded in 96 wells plates for imaging. One day later, cells were incubated at 37°C for 30 min with complete DMEM containing 5 µM Zinpyr-1 (green), Hoechst 33342 (blue) and 1 mM EDTA to chelate extracellular $Zn^{2+}$. After several washes in Live Cell Imaging Solution containing 1 mM EDTA, live cells were imaged with a CX7 high-content microscope and the mean average intensity per cell was quantified with the HCS Studio Cell Analysis software (n=3, >300 cells per experiment per condition). Statistical test: one-way ANOVA, *** p<0.001, **** p<0.0001. Scale bar: 30 µm.

**Supplementary Figure 10 – Tau oligomers uptake is unaffected by *C3orf58* or *SLC39A9* deficiency**

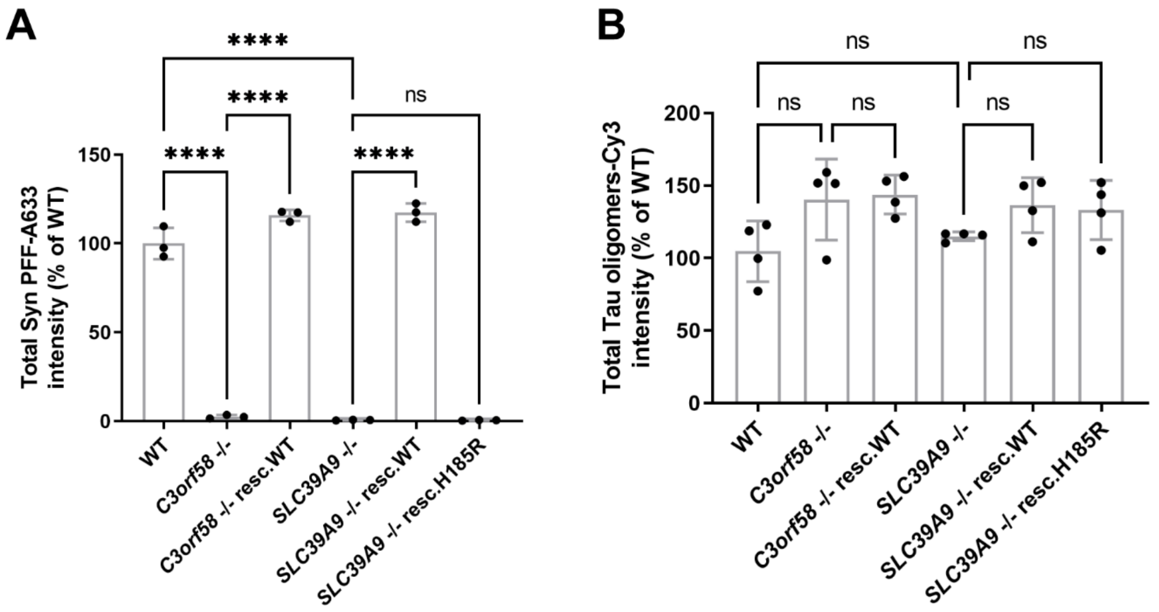


**(A)** WT RPE-1 cells were treated for 24 h with either 80 nM Tau oligomers-Cy3 (magenta), 60 nM α-syn-PFF-A633 (green) or both before fixation, nuclei staining with Hoechst (blue), and imaging using a CX7 high-content microscope. Scale bar: 30 µm. **(A,B)** WT, *C3orf58* -/-, *SLC39A9* -/- and indicated rescued lines were treated for 24 h with **(A)** 60 nM α-syn-PFF-A633 (green) or **(B)** 80 nM Tau oligomes-Cy3 (magenta). After fixation and nuclei staining with Hoechst (blue), cells were imaged using a CX7 high-content microscope and PFF-A633 or Tau oligomers-Cy3 mean total fluorescence intensity per cell were quantified with the HCS Studio Cell Analysis software. Data are shown as percent of WT values (mean ±sd). Statistical test: one-way ANOVA, * p<0.05, ** p<0.01, *** p<0.001, **** p<0.0001.

**Supplementary Figure 11 – Transcriptomic analysis of *C3orf58* $^{-/-}$ and *SLC39A9* $^{-/-}$ RPE-1 cells reveals highly similar changes in both lines, suggesting similar modes of action**

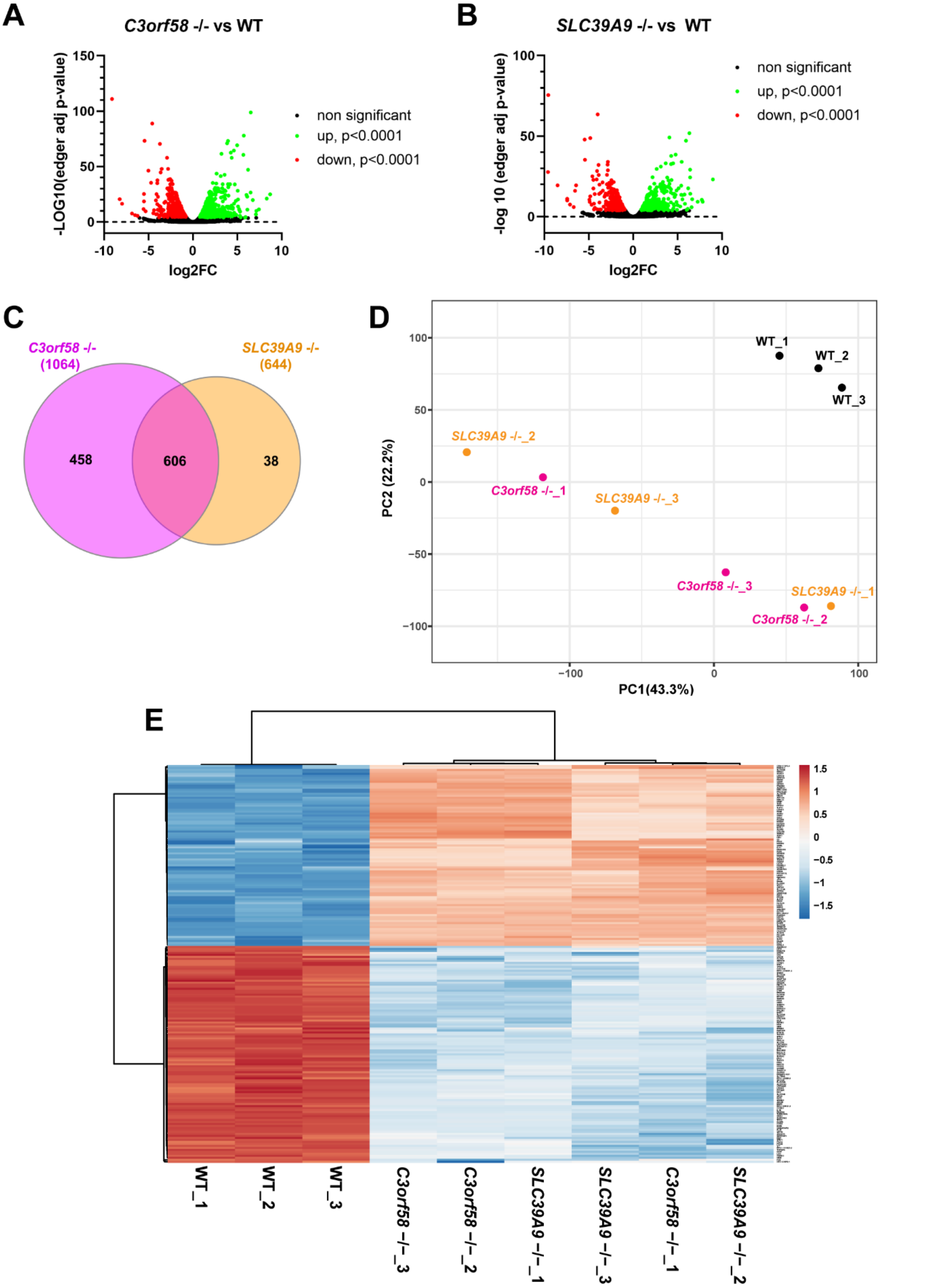


**(A,B)** RNAseq in *C3orf58* -/- and *SLC39A9* -/- RPE-1 cells followed by differential gene expression analysis compared to WT cells using the edgeR package (**A,** *C3orf58* -/- vs WT; **B**, *SLC39A9* -/- vs WT).

-LOG10 edgeR adjusted p-value are shown as a function of log2 foldchange. Upregulated and downregulated (p<0.0001) genes are shown as green and red dots, respectively. **(C)** Venn diagram showing the high number of differentially expressed genes (p<0.0001) that are common to *C3orf58* -/- and *SLC39A9* -/- cells. **(D)** PCA analysis of gene expression values in WT, *C3orf58* -/- and *SLC39A9* -/- cells, showing that both KO cluster separately from WT, but are not sufficiently different from each other to form distinct clusters. **(E)** Heatmap from clustering analysis of the 200 most differentially expressed genes from a contrast analysis between WT samples, and all KO samples of both genotypes together. Again, the clustering groups together all KO samples, regardless of the genotype, indicative of highly similar changes in both KO lines.

**Supplementary Figure 12 – KO of *C3orf58* or *SLC39A9* affect early uptake events**

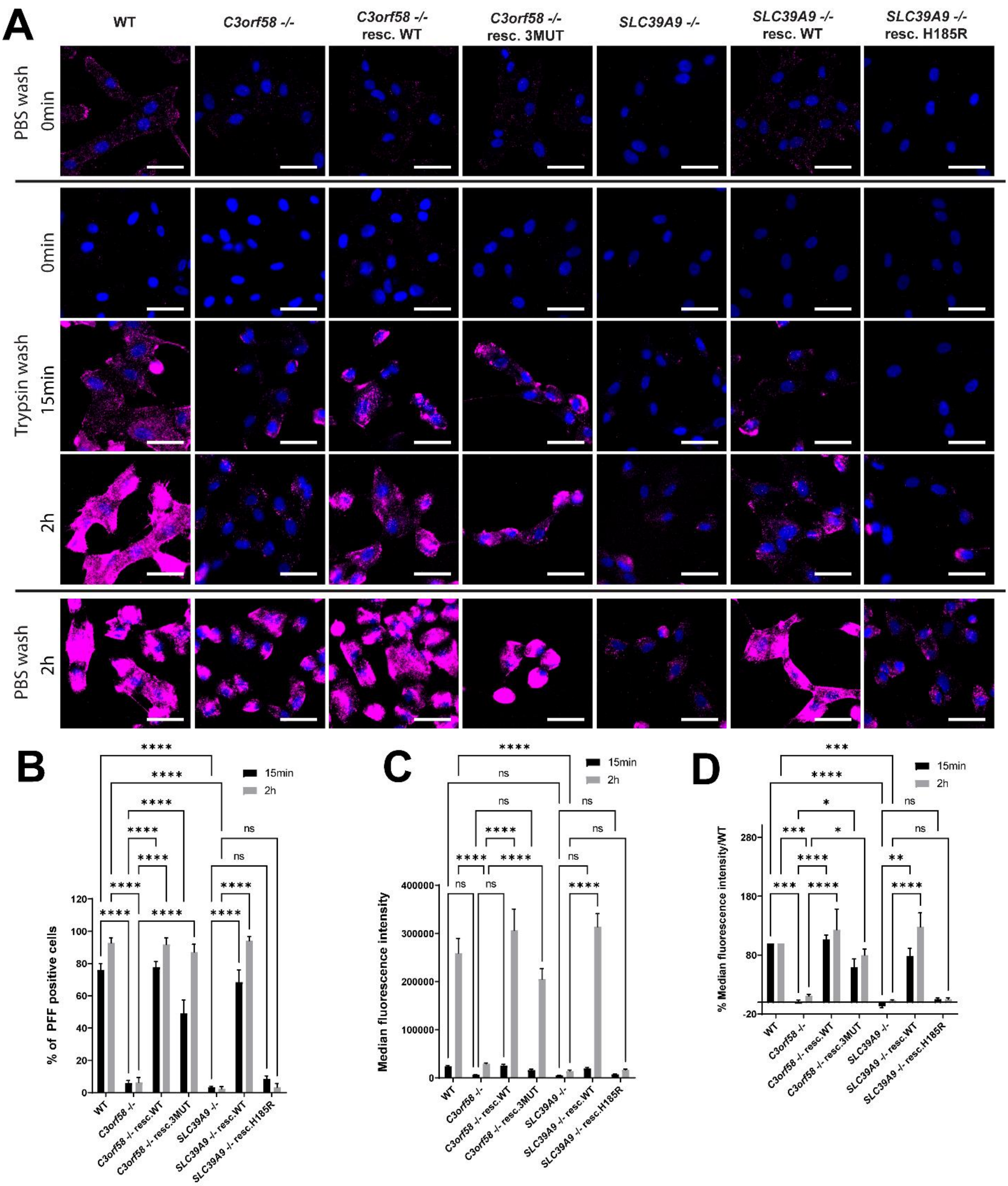


**(A)** RPE-1 cells of the indicated genotypes were incubated on ice with 180 nM α-syn-PFF-A633 for 20 min in serum-free DMEM. Cells were then incubated at 37°C for the indicated times, before treatment as follows: PBS wash (top and bottom panels) or 30 sec pre-warmed trypsin wash (middle panels, to remove surface-bound PFFs and retain only signal of internalized PFFs). Cells were then fixed, and nuclei stained with Hoechst 33342 before imaging on a Zeiss epifluorescence microscope at 20x. **(B,C,D)** The indicated values were quantified using ImageJ from the experiment in **(A)**: % of PFF positive cells **(B)**, median fluorescence intensity **(C)**, and relative median fluorescence intensity as % of control **(D)**. Statistical tests: 2way ANOVA; * p<0.05; ** p<0.01; *** p<0.001; **** p<0.0001. n=2 independent experiments.

**Supplementary Figure 13. Surface proteome analysis of *C3orf58*$^{-/-}$, *SLC39A9*$^{-/-}$, *TM9SF2*$^{-/-}$ and *VPS35*$^{+/-}$ RPE-1 cells**

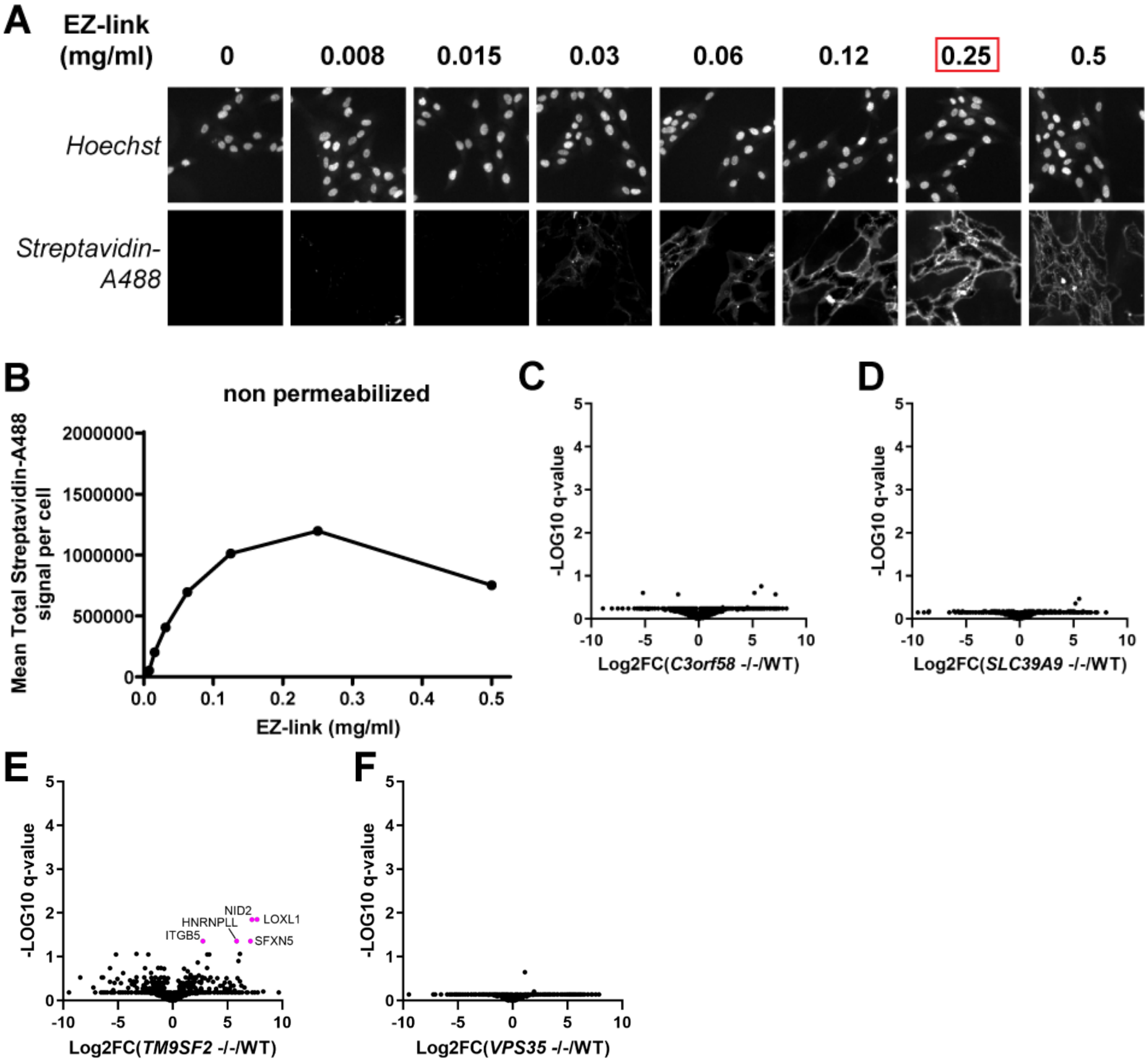


**(A)** RPE-1 cells were treated with increasing amounts of EZ-link biotinylation reagent for 30 min at 4°C (protocol adapted from Li et al.[49]). After quenching and fixation, nuclei were stained with Hoechst and biotinylated proteins with Alexa488-coupled Streptavidin. Images were acquired with a CX7 high-content microscope and streptavidin signals quantified with the HCS Studio Cell Analysis software (shown in **B**). 0.25 mg/ml provided optimal biotinylation of surface proteins, and this concentration was selected for scaling up. **(C,D)** Biotinylated surface proteins from WT, *C3orf58*$^{-/-}$, *SLC39A9*$^{-/-}$, *TM9SF2*$^{-/-}$ and *VPS35*$^{+/-}$ cells were subjected to streptavidin pulldown followed by LC-MS/MS analysis. Changes in protein levels (estimated by peptide counts) compared to WT cells are indicated as Log2 fold-change (**C**, *C3orf58*$^{-/-}$/WT; **D**, *SLC39A9*$^{-/-}$/WT; **E**, *TM9SF2*$^{-/-}$/WT; **E**, *VPS35*$^{+/-}$/WT). Significant changes were observed solely in *TM9SF2*$^{-/-}$ cells (multiparametric unpaired t-tests, FDR calculated with two-stage method for Benjamini, Krieger and Yekutieli).

**Supplementary Figure 14. C3orf58 is necessary for optimal colocalization of surface-bound PFFs with surface HS**

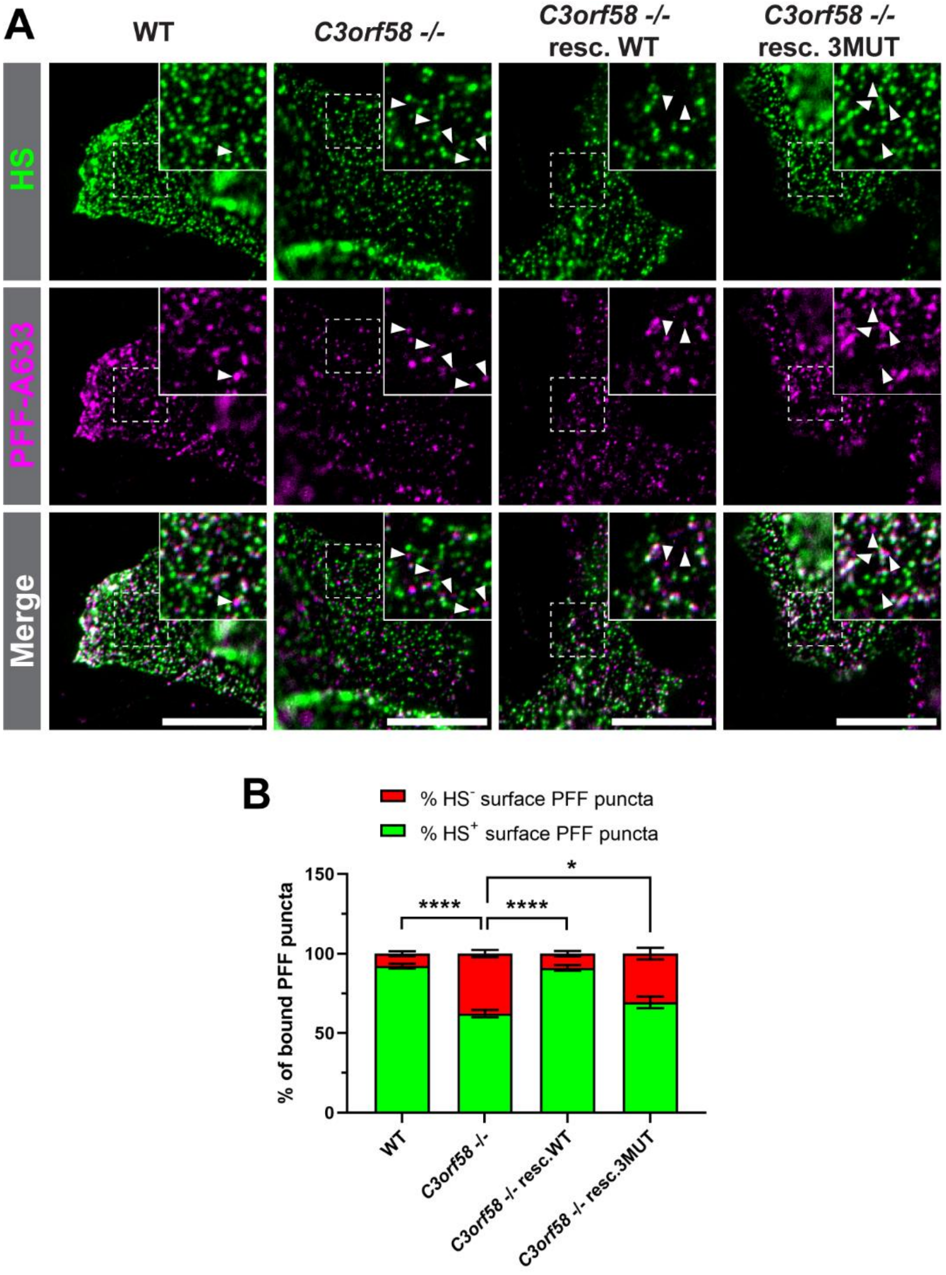


**(A)** PFF-A633 binding and surface HS immunostaining was performed on ice in live WT, *C3orf58* -/- or *SLC39A9* -/- monoclonal RPE-1 cells (± indicated rescue constructs), before fixation and nuclei staining with Hoechst. Images were obtained using a Zeiss fluorescence microscope at 63x. The insets show the punctate signals for HS and PFF at the cell surface. Scale bar: 20 µm. **(B)** The percentage of PFF puncta colocalizing with HS was quantified (n=3 independent experiments, >1000 dots quantified per condition). Kinase-dead C3orf58 only slightly rescued the reduced PFF/HS colocalization phenotype caused by C3orf58 deficiency, which might be due to either the importance of kinase activity, or slightly lower expression level and perturbed glycosylation of this mutant compared to WT C3orf58 (**Supplementary Figure 8C**). Statistical test: one-way ANOVA; ** p<0.01, *** p<0.001, ****p<0.0001.

**Supplementary Figure 15 – mRNA levels of glycosaminoglycan related genes is affected in *C3orf58* and *SLC39A9* KO cells**

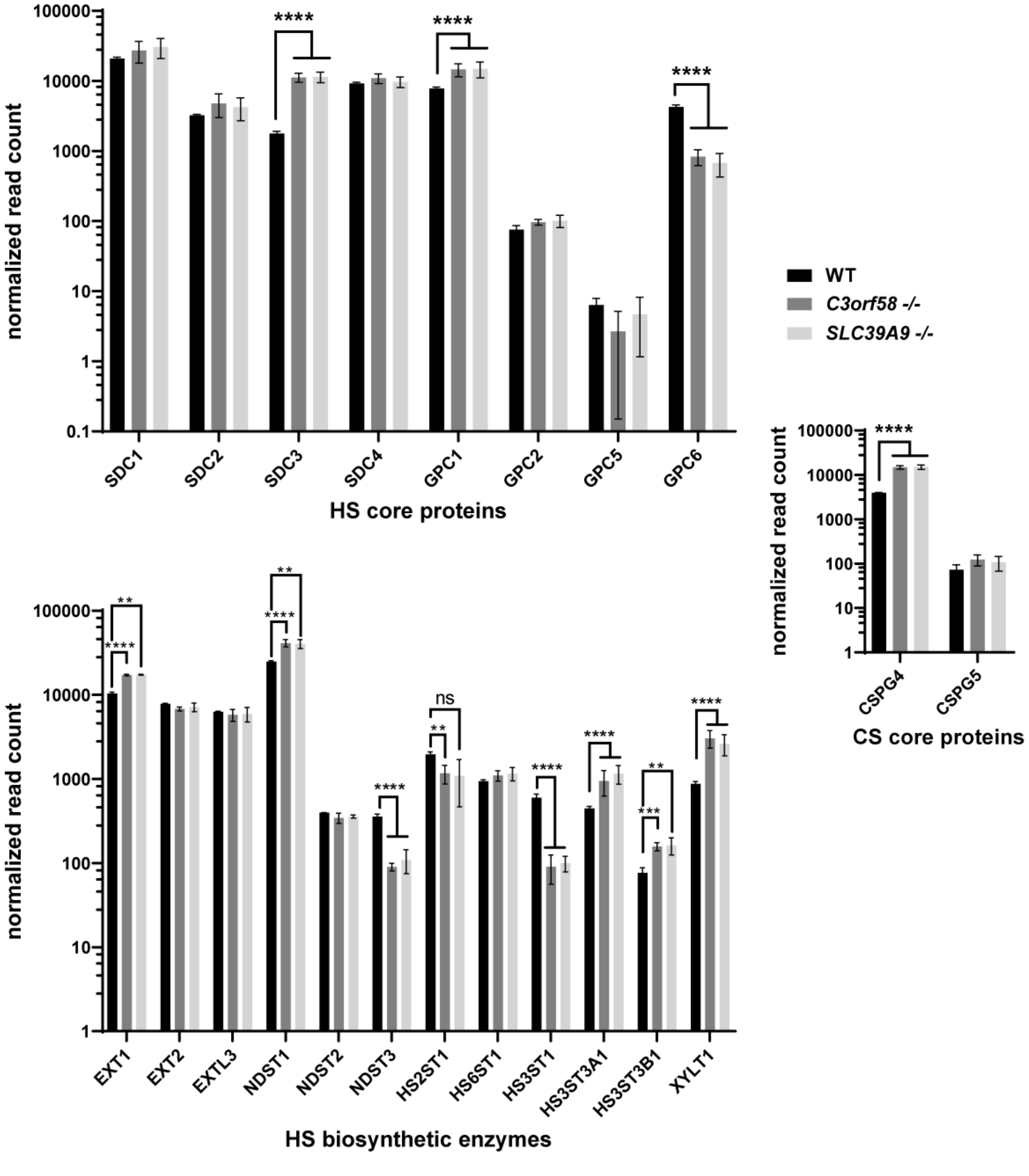


mRNA expression levels of the indicated groups of glycosaminoglycan related genes (top, HS core proteins; bottom, HS biosynthethic enzymes; right, chondroitin sulfate (CS) core proteins). Data are expressed as mean ±SD of normalized read counts (log scale), as determined by RNA sequencing in RPE-1 cells of the indicated genotypes (WT, black; *C3orf58* -/-, dark grey; *SLC39A9* -/-, light grey). Adjusted edger p-values were calculated with the edgeR package: ** adj. p<0.01; *** adj. p<0.001; **** adj. p<0.0001.

### Supplementary Figure 16. Changes in surface HS chains in *C3orf58*$^{-/-}$ and *SLC39A9*$^{-/-}$ RPE-1 cells

| Di/tetra-saccharide | Abbreviation | $R^1$ | $R^2$ | $R^3$ |
|---|---|---|---|---|
| △UA-GlcNAc | △IVA | -H | -H | -- |
| △UA2S-GlcNAc | △IIIA | -H | -$SO_3H$ | -- |
| △UA-GlcNAc6S | △IIA | -$SO_3H$ | -H | -- |
| △UA2S-GlcNAc6S | △IA | -$SO_3H$ | -$SO_3H$ | -- |
| △UA-GlcNS | △IVS | -H | -H | -- |
| △UA2S-GlcNS | △IIIS | -H | -$SO_3H$ | -- |
| △UA-GlcNS6S | △IIS | -$SO_3H$ | -H | -- |
| △UA2S-GlcNS6S | △IS | -$SO_3H$ | -$SO_3H$ | -- |
| △UA-GlcNAc6S-GlcA-GlcNS3S6S | Tetra-1 | -$SO_3H$ | -H | -Ac |
| △UA-GlcNS6S-GlcA-GlcNS3S6S | Tetra-2 | -$SO_3H$ | -H | -$SO_3H$ |
| △UA-GlcNS6S-IdoA2S-GlcNS3S6S | Tetra-3 | -$SO_3H$ | -$SO_3H$ | -- |
| △UA-GlcNS-IdoA2S-GlcNS3S | Tetra-4 | -H | -H | -- |
| △UA2S-GlcNS-IdoA2S-GlcNS3S | Tetra-5 | -H | -$SO_3H$ | -- |

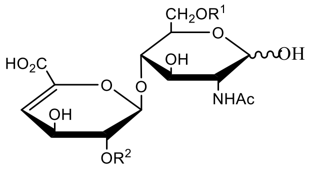


ΔIA-ΔIVA

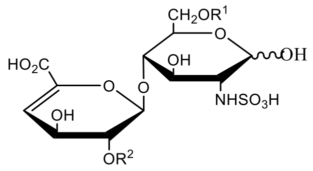


ΔIS-ΔIVS

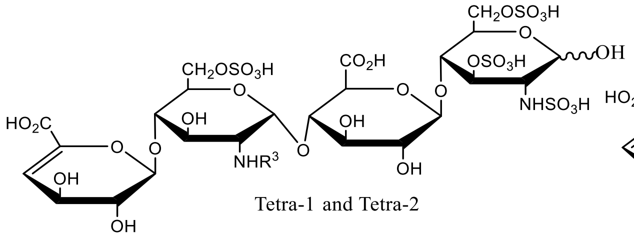


Tetra-1 and Tetra-2

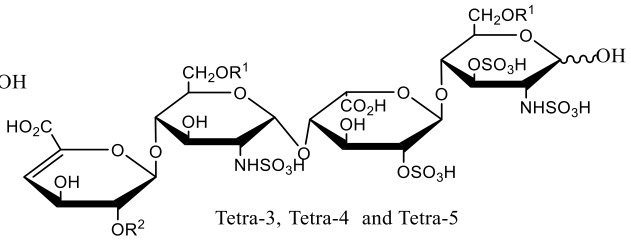


Tetra-3, Tetra-4 and Tetra-5

The HS disaccharides and tetrasaccharides analyzed in this study are described in the above table, together with the corresponding molecular structures. Complementary to Figure 7.

**Supplementary Figure 16. (continued)**

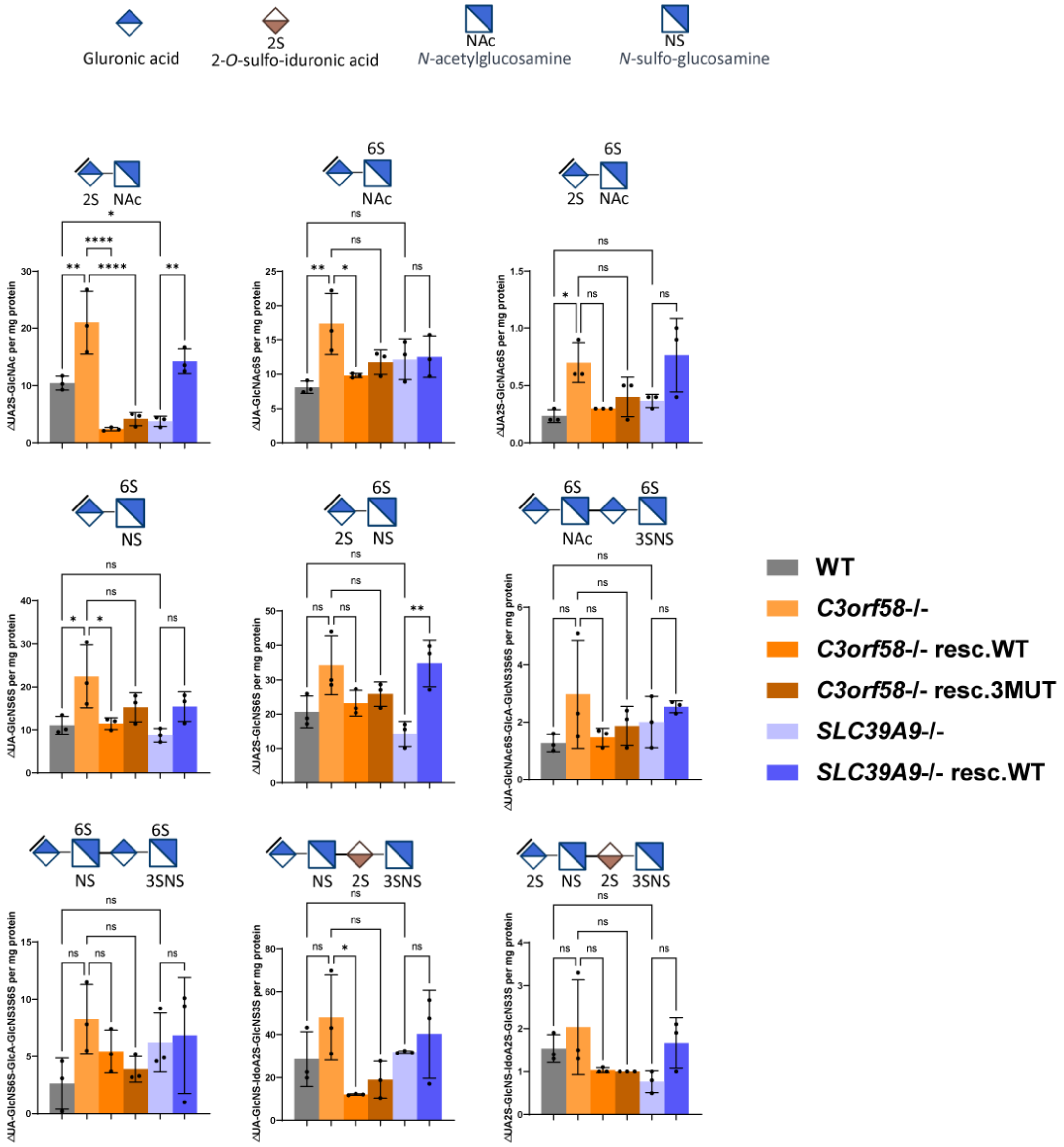


Bar graphs showing normalized quantities of the indicated HS disaccharides and tetrasaccharides isolated from the cell surface HS from WT, *C3orf58* or *SLC39A9* monoclonal KO RPE-1 cells (± indicated rescue constructs). LC-MS/MS with $^{13}$C-labeled internal standards was used. HS amounts were normalized to protein content from the cellular sample from which HS were isolated. A schematic representation of the analyzed species is added above each graph. Statistical tests: one-way ANOVA; * p<0.05, ** p<0.01, ****p<0.0001. Complementary to Figure 7.

### Supplementary Figure 16. (continued)

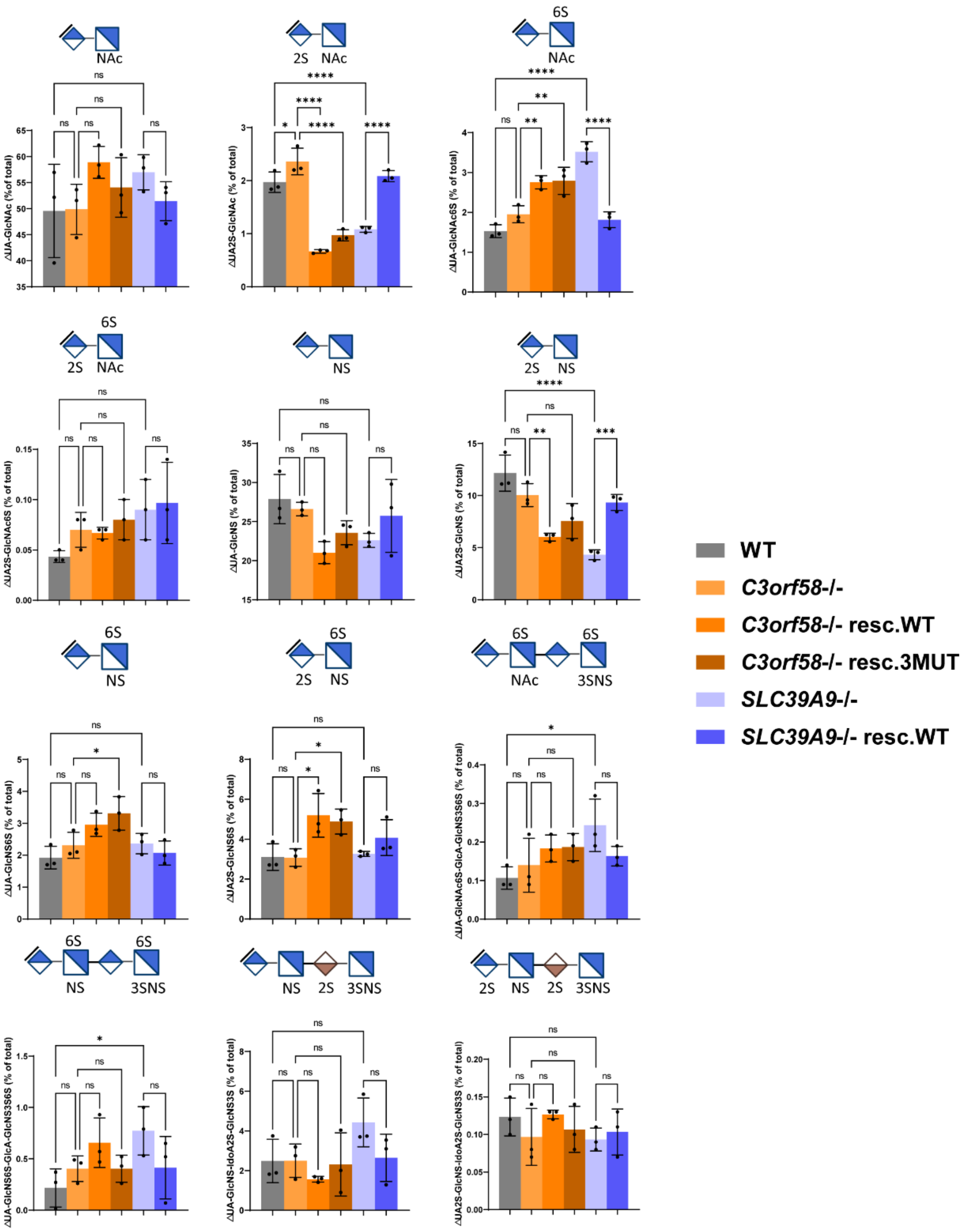


Surface HS composition (percentage of each disaccharide and tetrasaccharide species relative to total HS) of WT, *C3orf58* -/- or *SLC39A9* -/- monoclonal RPE-1 cells (± indicated rescue constructs). LC-MS/MS with $^{13}$C-labeled internal standards was used. A schematic representation of the analyzed species is added above each graph. Statistical tests: one-way ANOVA; * p<0.05, ** p<0.01, *** p<0.001, ****p<0.0001. Complementary to Figure 7.

Supplementary Figure 17. Golgi structure

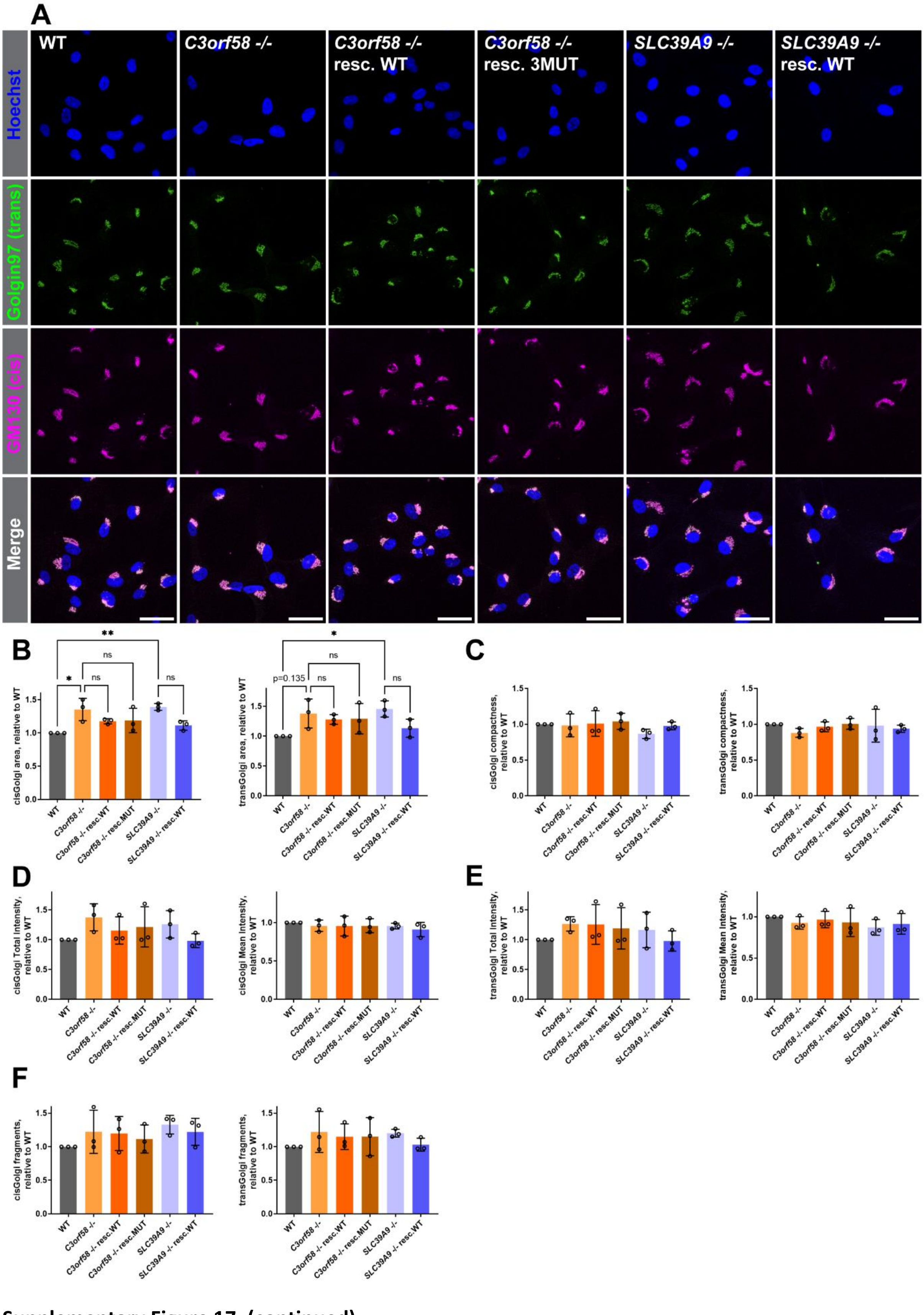


Supplementary Figure 17. (continued)

**(A)** RPE-1 cells of the indicated genotypes were fixed and stained by immunofluorescence for the cisGolgi marker GM130 (green) and the transGolgi marker Golgin 97 (magenta). Nuclei were stained with Hoechst. Max intensity projections of z-stacks acquired by confocal microscopy are shown. Scale bar: 50 µm. **(B-F)** A CellProfiler pipeline (adapted from[56]) was used to estimate cisGolgi and transGolgi morphology (**B**: area, **C**: compactness, **F**: number of fragments) and measure signal intensity (**D**: mean total intensity, **E**: mean average intensity) of both markers. Data are presented as mean ±SD. Statistical test: one-way ANOVA; * $p<0.05$, ** $p<0.01$.

**Supplementary Figure 18. Generation of *C3orf58* -/- and *SLC39A9* -/- iPSCs**

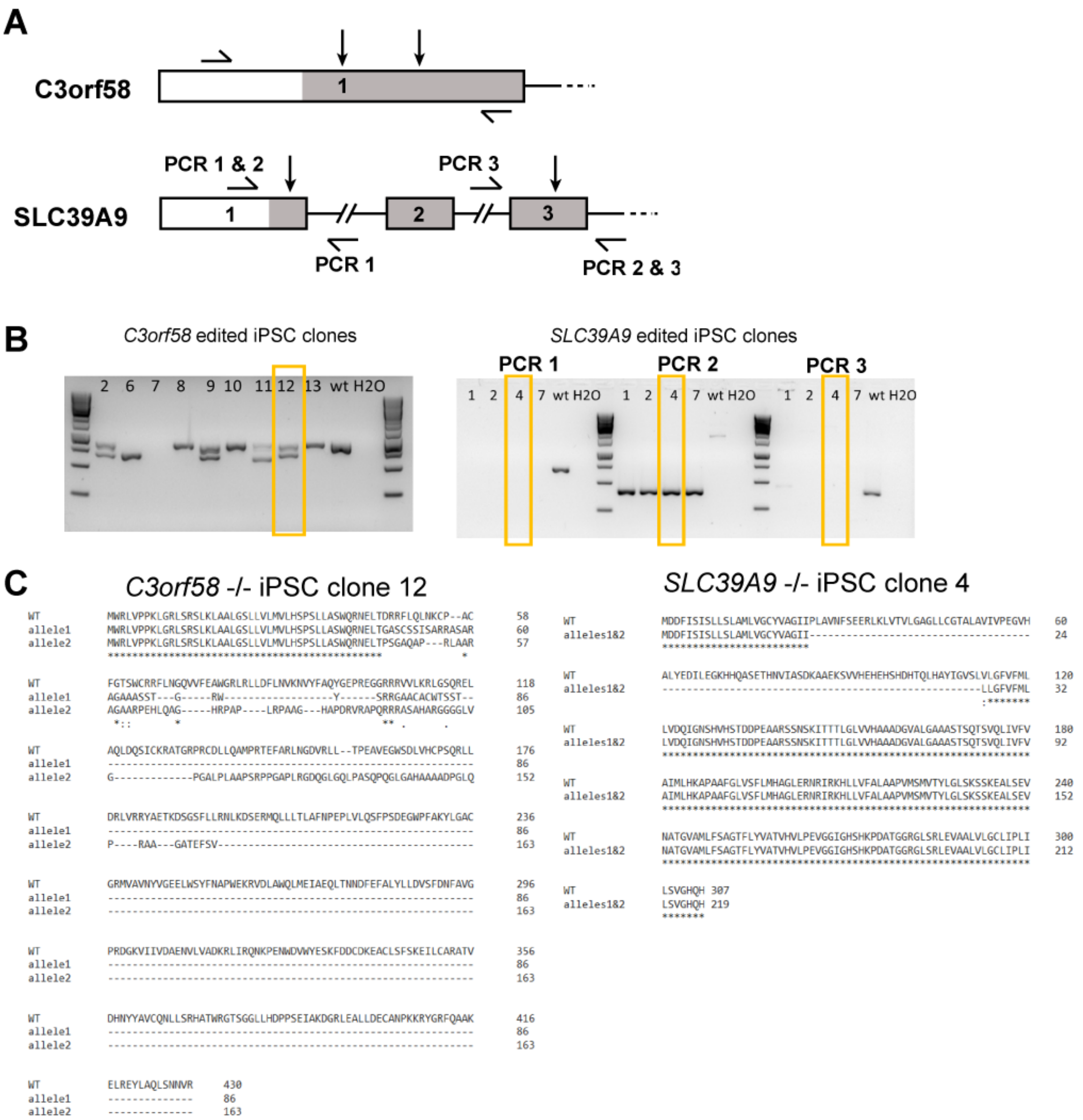


**(A)** Schematic representation of KO and PCR-screening strategies for the *C3orf58* and *SLC39A9* genes in human iPSCs. **(B)** Allelic sequences from the indicated CRISPR-edited iPSC clones were PCR-amplified using Q5 polymerase and oligonucleotides indicated in **(A)**. Orange boxes indicate selected clones. **(C)** PCR products from **(B)** were subjected to Sanger sequencing, and the obtained allelic sequences were translated *in silico* before alignment on the WT sequence. The identified allelic sequences are shown and aligned to the reference WT sequence. For *C3orf58* -/- clone 12, alleles 1 and 2 presented cumulated 20 bp (16 bp + 4 bp) or uninterrupted 179 bp deletions respectively, resulting in frameshift and premature stop codons. For *SLC39A9* -/- clone 4 iPSCs, a homozygous deletion (264 coding bp) resulted in loss of 88 amino acids from the remaining protein product (identical to allele 2 from *SLC39A9* -/- clone 10 RPE-1 cells used throughout the study, see Supplementary Figure 7C), which inactivates the $Zn^{2+}$ transporting function (Supplementary Figure 10).

**Supplementary Figure 18. (continued)**

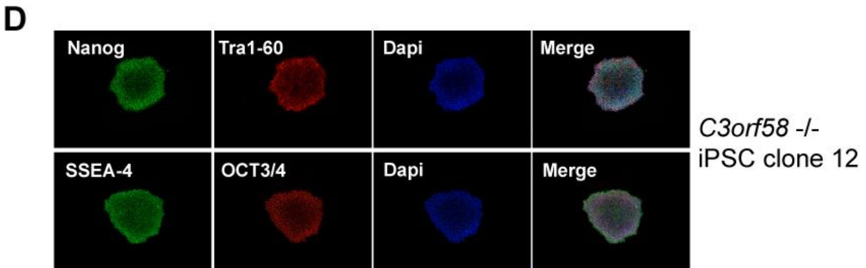

**(D)** Pluripotency of both edited iPSC lines was verified by immunostaining against the pluripotency markers Nanog, Tra1-60, SSEA-4 and OCT3/4. **(E)** Karyotype of *C3orf58* -/- and *SLC39A9* -/- monoclonal iPSCs. **(F)** CRISPR-edited iPSC showed a duplication of one copy of the chr20q region as assessed by a qPCR-based genome stability test, but this did not prevent proper differentiation into iMGL (see Supplementary Figure 19).

**Supplementary Figure 19. Human microglia derived from *C3orf58* and *SLC39A9* KO iPSC correctly express microglial markers**

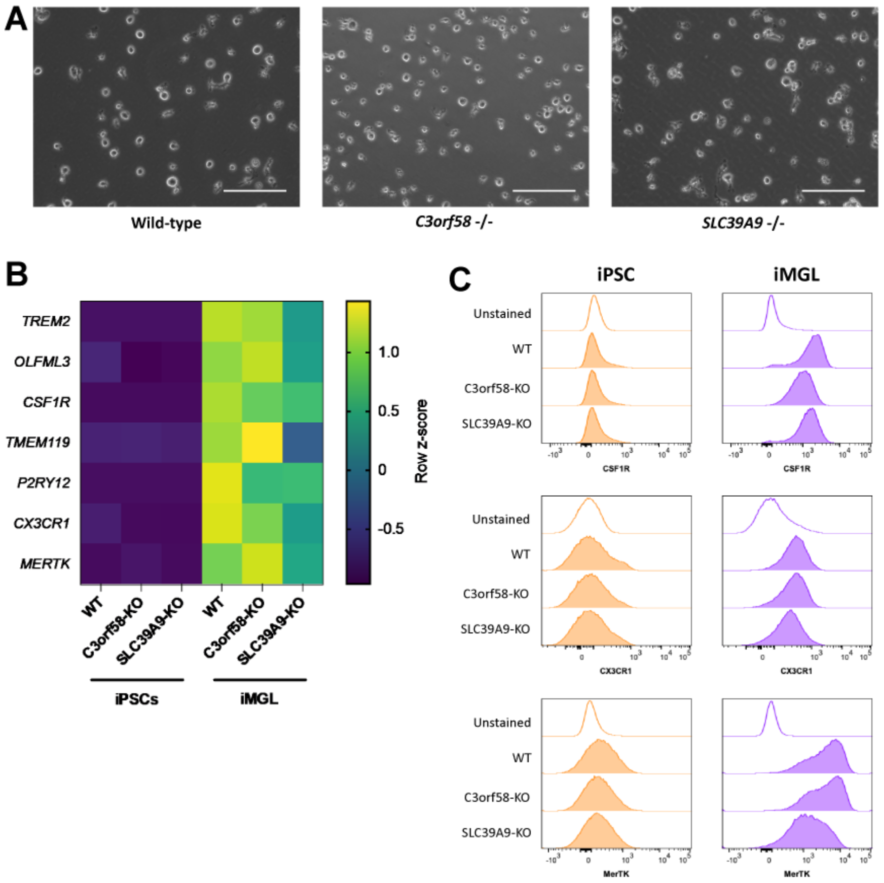


**(A)** Representative bright field images of WT, *C3orf58* -/- and *SLC39A9* -/- human iPSC-derived microglia (iMGL) used in the study. Scale bar = 150 µm. **(B)** qRT-PCR was used to measure mRNA levels of the indicated microglial marker genes in iPSCs or iMGL of the indicated genotypes. Levels were normalized to *GAPDH* and *YWHAZ* genes. iPSC n=4, iMGL n=3. **(C)** Surface microglial markers CSF1R, CX3CR1 and MerTK expression levels were assessed by flow cytometry in iPSC or iMGL of the indicated genotypes.

**Supplementary Figure 20 – α-syn PFF uptake is partially mediated by macropinocytosis in RPE-1 cells**

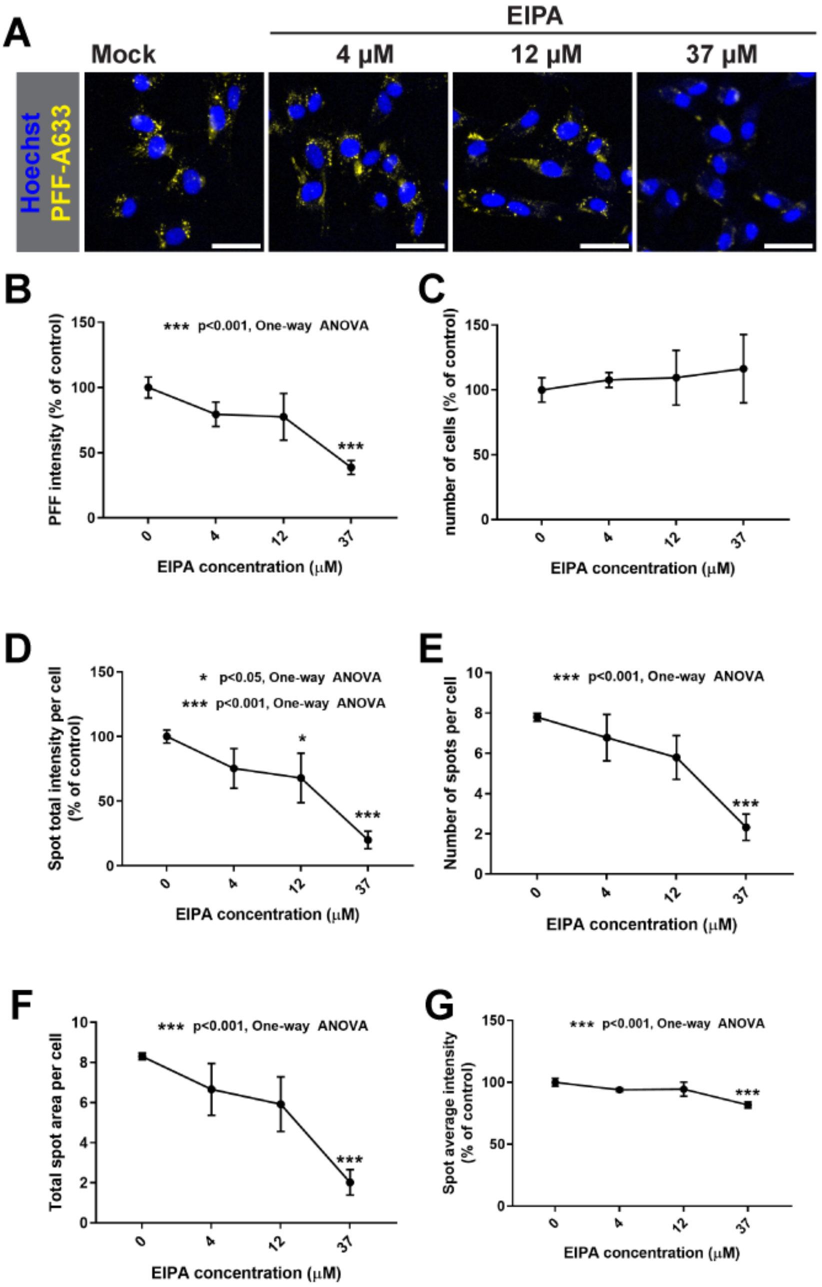


**(A)** RPE-1 cells were treated with increasing concentrations of the macropinocytosis inhibitor EIPA for 30 min at 37°C before 14h overnight treatment with 60 nM PFF-A633, fixation, nuclei staining with Hoechst, imaging and **(B-G)** quantification with a CX7 high-content microscope. Parameters quantified are shown as mean ±SD: **(B)** total PFF intensity, **(C)** cell number, **(D)** total PFF spots intensity per cell, **(E)** number of PFF spots per cell, **(F)** area covered by PFF spots per cell, and **(G)** average intensity within PFF spots. 37 μM treatment resulted in significant decrease in PFF uptake.
